# Coil-to-Helix Transition at the Nup358-BicD2 Interface Activates BicD2 for Dynein Recruitment

**DOI:** 10.1101/2021.05.06.443034

**Authors:** James M. Gibson, Heying Cui, M. Yusuf Ali, Xiaoxin Zhao, Erik W. Debler, Jing Zhao, Kathleen M. Trybus, Sozanne R. Solmaz, Chunyu Wang

**Affiliations:** Department of Biological Sciences, Department of Chemistry and Chemical Biology, Center for Biotechnology and Interdisciplinary Studies, Rensselaer Polytechnic Institute, 110 8^th^ Street, Troy NY 12180.; Department of Chemistry, Binghamton University, P.O. Box 6000, Binghamton, NY 13902.; Department of Molecular Physiology and Biophysics, University of Vermont, Burlington, Vermont.; Department of Biochemistry & Molecular Biology, Thomas Jefferson University, 1020 Locust Street, Philadelphia, PA 19107.

## Abstract

Nup358, a nuclear pore protein, facilitates a nuclear positioning pathway that is essential for many biological processes, including neuromuscular and brain development. Nup358 binds and activates the auto-inhibited dynein adaptor Bicaudal D2 (BicD2), which in turn recruits and activates the dynein machinery to position the nucleus. However, the molecular details of the Nup358/BicD2 interaction remain poorly understood. Here, we show that a minimal Nup358 domain activates dynein/dynactin/BicD2 for processive motility on microtubules. Using nuclear magnetic resonance (NMR) titration and chemical exchange saturation transfer (CEST), a Nup358 helix encompassing residues 2162-2184 was identified, which transitioned from random coil to an α-helix upon BicD2-binding and formed the core of the Nup358-BicD2 interface. Mutations in this region of Nup358 decreased the Nup358/BicD2 interaction, resulting in decreased dynein recruitment and impaired motility. BicD2 thus recognizes the cargo adaptor Nup358 though a “cargo recognition α-helix”, a structural feature that may stabilize BicD2 in its activated state and promote processive dynein motility.

## Introduction

Cytoplasmic dynein is the predominant motor responsible for minus-end directed traffic on microtubules^1^, which facilitates a vast number of transport events that are critical for chromosome segregation, signal transmission at synapses, and brain and muscle development^2–16^. Integral to the transport machinery are *dynein adaptors*, such as Bicaudal D2 (BicD2), whose N-terminal region (BicD2^CC1^) recruits and activates dynein-dynactin for processive motility ^17–27^. Also integral to the dynein transport machinery are *cargo adaptors*, which bind to the C-terminal domain (CTD) of BicD2^28^. Cargo-adaptors are required to activate BicD2 for dynein binding, which is a key regulatory step for dynein-dependent transport^17–27^. In the absence of cargo adaptor/dynein adaptor complexes such as Nup358/BicD2, dynein and dynactin are auto-inhibited and only show diffusive motion on microtubules. Furthermore, in the absence of cargo adaptors, BicD2 assumes a looped, auto-inhibited conformation, in which its N-terminal dynein/dynactin binding site binds to the CTD and remains inaccessible. The CTD is required for auto-inhibition, as a truncated BicD2 without the CTD activates dynein/dynactin for processive motility. Binding of dynein adaptors/cargo to the CTD releases auto-inhibition, likely resulting in an extended conformation that recruits dynein and dynactin^17–27^. The crystal structures of the C-terminal cargo recognition domains of three BicD2 homologs have been determined^26,29,30^; However, the structural mechanisms of BicD2-mediated cargo recognition and dynein activation still remain poorly understood.

The cargo adaptors for human BicD2 include Nup358^11^, Rab6^GTP 28^ and nesprin 2-G^6^. Nup358, also known as RanBP2, is a 358 kDa nuclear pore complex protein with multiple functions^31^. During G2 phase, Nup358 engages in a pathway for positioning of the nucleus relative to the centrosome along microtubules by binding to BicD2, which in turn recruits dynein and dynactin (Fig. 1)^11^. This pathway is essential for apical nuclear migration during differentiation of radial glial progenitor cells, which give rise to the majority of neurons and glia cells of the neocortex ^4,5^. A second nuclear positioning pathway is facilitated by BicD2/dynein and nesprin 2-G^6^, a component of LINC (Linker of Nucleoskeleton and Cytoskeleton) complexes (LINC)^32^, which is important for migration of post-mitotic neurons during brain development^6^. Finally, Rab6^GTP^ recruits BicD2/dynein for the transport of Golgi-derived secretory vesicles^28,33^. Thus, BicD2 plays important roles in faithful chromosome segregation, neurotransmission at synapses as well as brain and muscle development ^4–6,8,11,28^. Mutations in BicD2 cause neuromuscular diseases including a subset of spinal muscular atrophy^34–38^, the most common genetic cause of infant death^39^.

**Fig. 1.**
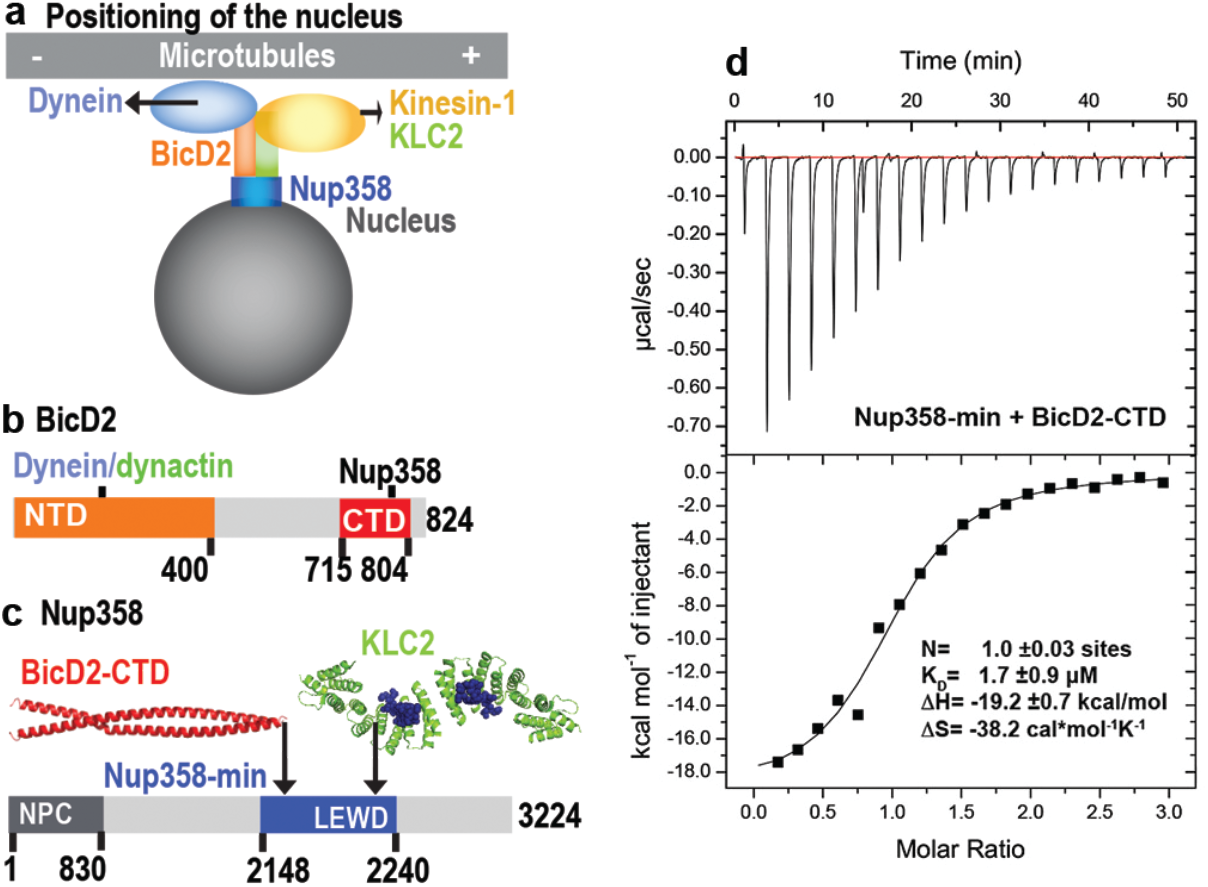
A minimized Nup358 domain interacts with BicD2 with micromolar affinity. (a) Nup358 interacts with both BicD2/dynein/dynactin and kinesin-1 (via kinesin-1 light chain 2, KLC2) to mediate bi-directional nuclear positioning in G2 phase of the cell cycle^11,46,47^. This pathway is essential for a fundamental process in brain development that is required for radial glial progenitor cells to differentiate to the majority of neurons and glia cells of the neocortex^4^. (b, c) Schematic representation of the expression constructs BicD2-CTD (b, red) and Nup358-min (c, blue), in the context of the full-length proteins (grey). (c) KLC2 is recruited to Nup358-min via a W-acidic motif with the sequence LEWD^47^. The X-ray structures of the TPR domain of the KLC2 (green, referred to as KLC2 hereafter), fused to a LEWD binding motif (purple)^49^ and the X-ray structure of the BicD2-CTD (red)^30^ are shown in cartoon representation in c. The α-helical N-terminal domain of Nup358 (dark grey) promotes anchorage into the NPC (d) A representative ITC thermogram of Nup358-min and BicD2-CTD is shown, from which the affinity (dissociation constant K_D_) was determined to be 1.7 ± 0.9 μM. N: number of sites; ΔS: the change in entropy; ΔH: the change in enthalpy. The experiment was repeated three times. The error of K_D_ was calculated as the standard deviation from three experiments. In Fig. S1, the affinity of Nup358-min-GST (i.e., with the GST-tag intact) and BicD2-CTD was determined by ITC to be 1.6 ± 1.0 μM.

Within the Nup358 sequence, there are many intrinsically disordered regions (IDR), commonly found in dynein adaptors, cargo adaptors, dynein and other factors involved in dynein-dependent transport^10,40–43^. Although IDRs and IDPs (intrinsically disordered proteins) make up ~30% of eukaryotic proteins and have important physiological functions^44^, they remain the most poorly characterized class of protein in their structure, dynamics and interactions. An example of IDR is a region in the dynein light intermediate chain 1 (LIC1), which forms a 15-residue α-helix to interact with the N-terminal domain of BicD2, an important step in the activation of dynein for processive motility ^40,41,43^. In addition to the interaction between LIC1 and BicD2, a larger interface is formed between the N-terminal coiled coil of BicD2, the dynein tail and dynactin, which promotes activation of dynein for processive motility^17,18,21,22^. Because of the small binding domain of LIC1 located in an IDR, this type of interactions can in some cases associate and dissociate at a quick rate while maintaining a high degree of specificity and affinity^45^, making them uniquely suited for transport processes.

In addition to BicD2, Nup358 also recruits the opposite polarity motor kinesin-1 via the subunit kinesin-1 light chain 2 (KLC2) which binds to a W-acidic motif with the sequence LEWD in Nup358^46–48^. While dynein is the predominant motor in G2 phase, kinesin-1 is also actively involved in nuclear positioning in G2 phase, modulating overall motility^11^. Such bidirectional transport is also displayed by mitochondria, endosomes, viruses, phagosomes, secretory vesicles, and many vesicles in neuronal axons and growth cones^2,3,6,9–16^. Opposite polarity motors often bind in close spatial proximity to cargo adaptors, but it is unknown how their overall motility is regulated.

Here we have determined the structural properties of the interface of a minimal Nup358/BicD2 complex by an interdisciplinary approach that combines NMR spectroscopy, X-ray crystallography, mutagenesis, circular dichroism spectroscopy and small-angle X-ray scattering. Importantly, this work is further enhanced by single-molecule binding and processivity assays which confirm the results obtained from the minimal complex in the context of intact dynein/dynactin/BicD2 motors and provide mechanistic insights into dynein activation. These results establish a structural basis for cargo recognition and suggest that Nup358 interacts with BicD2 through a “cargo recognition” α-helix. Single-molecule assays show that a minimal dimerized Nup358 construct is sufficient to activate dynein/dynactin/BicD2 for processive motility, and single molecule binding assays show that dynein-dynactin enhances the formation of the Nup358/BicD2 complex, thus providing mechanistic insights into dynein activation. Intriguingly, our NMR data also show that the binding site of BicD2 in Nup358 is spatially close to but does not overlap with the LEWD motif that acts as a kinesin-1 light chain 2 (KLC2) binding site, suggesting that kinesin and dynein machineries may interact simultaneously via Nup358.

## Results

### ITC establishes a minimal complex for Nup358/BicD2 interaction

Previously, we have determined an X-ray structure of the C-terminal domain of human BicD2 (BicD2-CTD, residues 715-804), which contains the binding sites for cargoes, including human Nup358 (Fig. 1b)^30^. A complex was reconstituted with BicD2-CTD and a minimal fragment of human Nup358 containing residues 2148-2240, which is called Nup358-min^30,47^ (Fig. 1c). Here, the affinity of the BicD2-CTD towards Nup358-min was determined by isothermal titration calorimetry (ITC) (Fig. 1d). The ITC thermogram fits well to a one-site binding model (number of sites N =1.0) and is thus consistent with a molar ratio of [Nup358]/[BicD2] of 1. This molar ratio is in agreement with our previously published molar masses obtained from size exclusion chromatography coupled to multi-angle light scattering (SEC-MALS), which showed that Nup358 and BicD2 form a 2:2 complex^50^. The one-site binding model is in line with a Nup358-dimer binding to a single binding site on a BicD2 dimer, although we cannot exclude the possibility that two Nup358 monomers bind to two binding sites on BicD2, where both sites have the same affinity. The equilibrium dissociation constant K_D_ was determined to be 1.7 ± 0.9 μM, in a similar range as observed for other BicD2/cargo complexes^50^. This is similar to the previously published affinity of 0.4 μM, obtained for BicD2-CTD towards a larger fragment of Nup358 (residues 2006-2443, with the GST-tag intact)^50^. To ensure the cleavage of GST-tag did not affect the binding affinity, we also performed an ITC-titration with Nup358-min-GST (i.e., with the GST-tag intact, whereas the GST was cleaved off in the first experiment) and the BicD2-CTD which yielded a very similar affinity of 1.6 ± 1.0 μM (Fig. S1). These ITC results confirm the mapped boundaries of the minimal binding site. The change of enthalpy was −19.2 ± 0.7 kcal/mol and the change of entropy –38.2 ± 5.8 cal/mol/K. Thus, the Nup358-BicD2 interaction is driven by a favorable enthalpy change, which overcomes the unfavorable entropy change.

### A dynein-dynactin-BicD2-Nup358^min^ complex moves processively

Single molecule reconstitutions were used to determine if Nup358^min^ can bind and relieve BicD2 auto-inhibition, which in turn allows dynein-dynactin to be recruited and activated for processive motion. We first reconstituted a dynein-dynactin-BicD2-Nup358^min^ (DDBN^min^) complex in which Nup358^min^ and full-length BicD2 (BicD2) were labeled with two different color Quantum dots (Qdots) and mixed with tissue purified unlabeled dynein-dynactin (DD) complex (Fig. 2a, but without microtubules). Only 24% dual-color complexes were observed for dynein-dynactin-BicD2-Nup^min^ (DDBN^min^) complex (Fig. 2b, d). We next reconstituted the complex with a Nup358^min^ that was dimerized with a leucine-zipper (hereafter called Nup358^min-zip^). Nup358^min^ dimerization enhanced the number of dual-colored DDBN complexes to 36% (Fig. 2b, c). The rationale for this strategy was based on our previous observation that dimerization of the *Drosophila* mRNA binding adaptor protein Egalitarian with a leucine zipper enhanced its affinity for BicD and bypassed the requirement for mRNA cargo for BicD activation^24^. All further single molecule reconstitutions used the dimerized version of Nup358-min (Nup358^min-zip^) because of its enhanced affinity for BicD2.

**Fig. 2.**
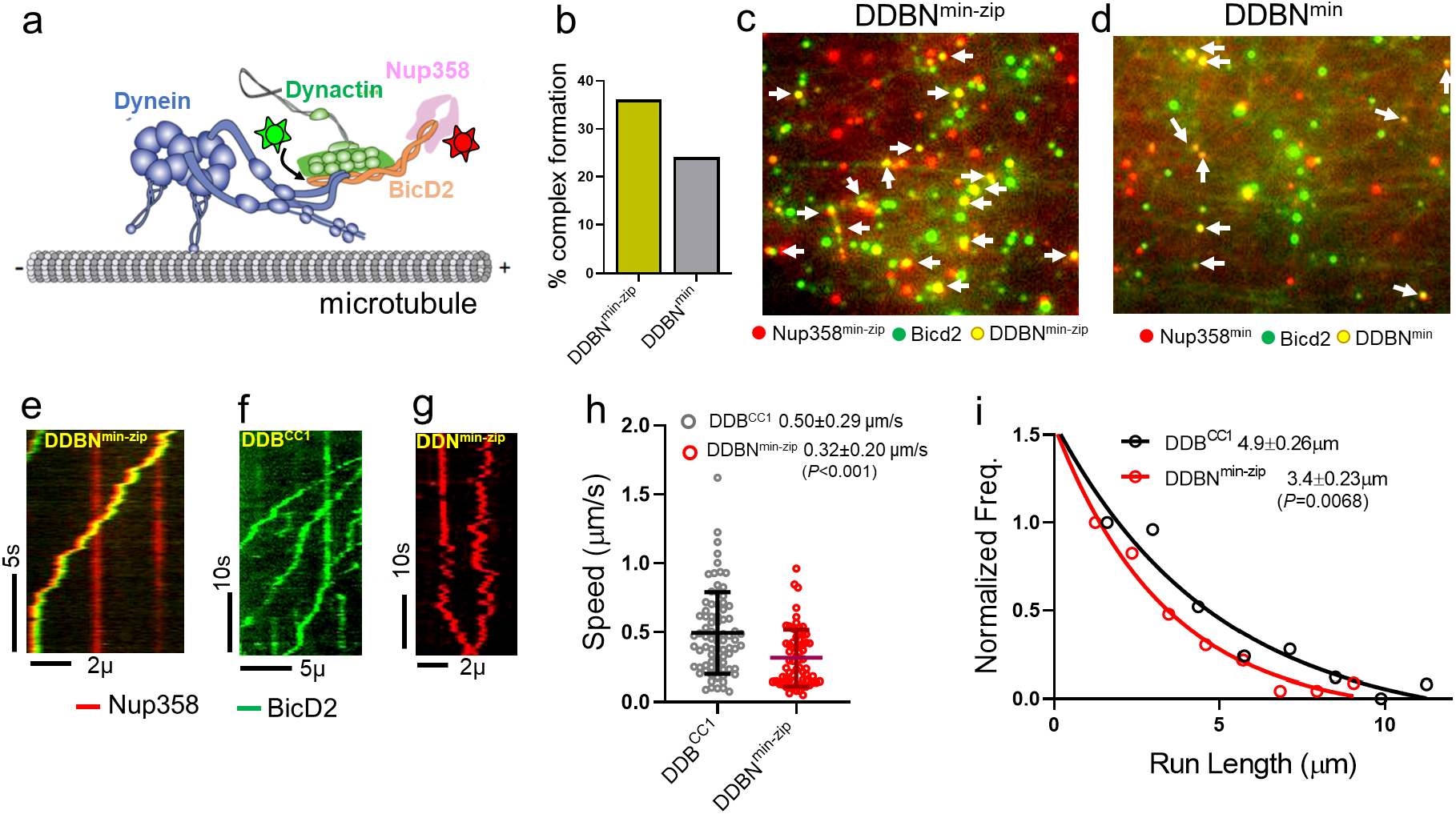
Nup358^min-zip^ is capable of forming a dynein-dynactin-BicD2-Nup358^min-zip^ complex (DDBN^min-zip^) that is activated for processive motility. (a) Schematic of a DDBN^min-zip^ complex bound to a microtubule. BicD2 and Nup358 are shown labeled with two different color Quantum dots (stars). (b) Dimerization of Nup358^min^ increases the formation of DDBN^min-zip^. A leucine-zippered Nup358 (Nup358^min-zip^) forms DDBN complex (yellow) at a higher frequency than that of Nup358 monomer (Nup358^min^) (grey). The percent formation of DDBN^min-zip^ complex was (36.1%, n=945) 50% higher than that of DDBN^min^ complexes (24.1%, n=514). (c, d) Visualization of complex formation of DDBN formed with a dimerized Nup358^min-zip^ versus a monomeric Nup358^min^. Yellow dots (indicated by white arrows) show the formation of the DDBN complex. Single color Qdot green indicates BicD2, and red indicates Nup358^min-zip^. (e) Kymograph of the DDBN^min-zip^ complex moving processively on microtubules, with Nup358^min-zip^ labeled with a 655 nm Qdot and BicD2 labeled with a 565 nm Qdot. (f) As a control, motion of dynein-dynactin in the presence of the N-terminal domain of BicD2 (BicD2^CC1^), with BicD2^CC1^ labeled with a 565 nm Qdot. (g) Dynein-dynactin and Nup358 complexes (DDN^min-zip^) showed diffusive movement on microtubules, with Nup358^min-zip^ labeled with a 655 nm Qdot. (h, i) Speed and run length of DDBN^min-zip^ (red) were compared with the constitutively active complex DDB^CC1^. The speed and run length of DDB^CC1^ are 0.50±0.29 μm/s (N=80) and 4.9±0.26 μm (N=80), compared with values of 0.32±0.20 μm/s (N=68) and 3.4±0.23 μm (N=68) for DDBN^min-zip^. Speeds for the two complexes are significantly different (p<0.001, Unpaired t-test with Welch’s correction), as are the run lengths (p=0.0068, Kolmogorov-Smirnov test).

The motion of the dynein-dynactin-BicD2-Nup358^min-zip^ (DDBN^min-zip^) complex on microtubules (Fig. 2a) was quantified. The DDBN^min-zip^ complex exhibited robust processive motion on surface-immobilized MTs, implying that Nup358^min-zip^ relieves BicD2 auto-inhibition to allow dynein activation (Fig. 2e, Movie S1). The speed and run-length of all dual-color DDBN^min-zip^ complexes were analyzed, with the run length obtained from a one-phase exponential decay fit, and speed determined from the Gaussian distribution fit. The speed and run-length of DDBN^min-zip^ were quantified as 0.32±0.20 μm/s (n=68) and 3.4±0.23 μm (n=68), respectively (Fig. 2h, i). These motile properties of DDBN^min-zip^ were compared with that of the DDB^CC1^ complex (containing the BicD2 N-terminal coiled-coil 1 (CC1) domain), which has been well established to be a fully active complex^21,24^ (Fig. 2f). The speed and run-length of the DDB^CC1^ complex was 0.50±0.29 μm/s (n=80) and 4.9±0.13 μm (n=80), respectively, which are significantly faster (p<0.001) and longer (p=0.0068) than that of DDBN^min-zip^(Fig.2h, i). Nonetheless, the directed motion of DDBN^min-zip^ is very different from the auto-inhibited dynein-dynactin complex, which shows only diffusive movement on MTs^24^.

To determine whether Nup358^min-zip^ can also bind directly to dynein-dynactin in the absence of BicD2, we formed a complex of dynein-dynactin-Nup358^min-zip^ (DDN^min-zip^) in which Nup358^min-zip^ was labeled with 655nm Qdots. This DDN^min-zip^ complex binds to MTs through dynein, but exhibits diffusive motion on MTs (Fig. 2g; Movie S2), implying that both BicD2 and Nup358 are required for dynein-dynactin processive motion.

As a control, we confirmed that unlike the DDN^min-zip^ complex, Nup358^min-zip^ alone does not bind to microtubules (data not shown). Thus, we conclude that Nup358^min-zip^ interacts directly with dynein/dynactin. Our results suggest that Nup358 relieves the auto-inhibition of BicD2 so that dynein-dynactin can bind and move processively on MTs.

### NMR titration narrowed the BicD2 binding site to the N-terminal half of Nup358-min

Because our single molecule processivity assays confirmed that the Nup358-min domain forms a DDBN complex that is activated for processive motility, we characterized the BicD2-binding sites on Nup358-min, employing solution NMR, which can provide atomic resolution information for protein interactions in the native solution state. First, backbone assignment of Nup358-min was carried out using standard triple resonance experiments, 3D HNCO^51^, HNCACO^52^, HNCA^51^, HNCACB^53^, and CBCACONH^54^. 82 out of the 89 possible backbone amides were assigned in the ^1^H-^15^N HSQC (Fig. S2). Then ^15^N labeled Nup358-min was titrated with unlabeled BicD2-CTD. Fig. 3a shows the overlay of the HSQC spectrum of apo Nup358-min with that of the complex, for which ^15^N labeled Nup358-min and unlabeled BicD2-CTD were mixed at a 1:1 molar ratio. The HSQC spectrum of the apo Nup358-min is characterized by a lack of dispersion, consistent with an intrinsically disordered protein (IDP). Although the addition of BicD2-CTD resulted in little peak movement in the HSQC, significant peak intensity changes were observed for the N-terminal half of Nup358-min (Fig. 3a and 3b). This points to a relatively wide binding region undergoing intermediate to slow exchange on the NMR time scale, due to ligand binding.

**Fig. 3.**
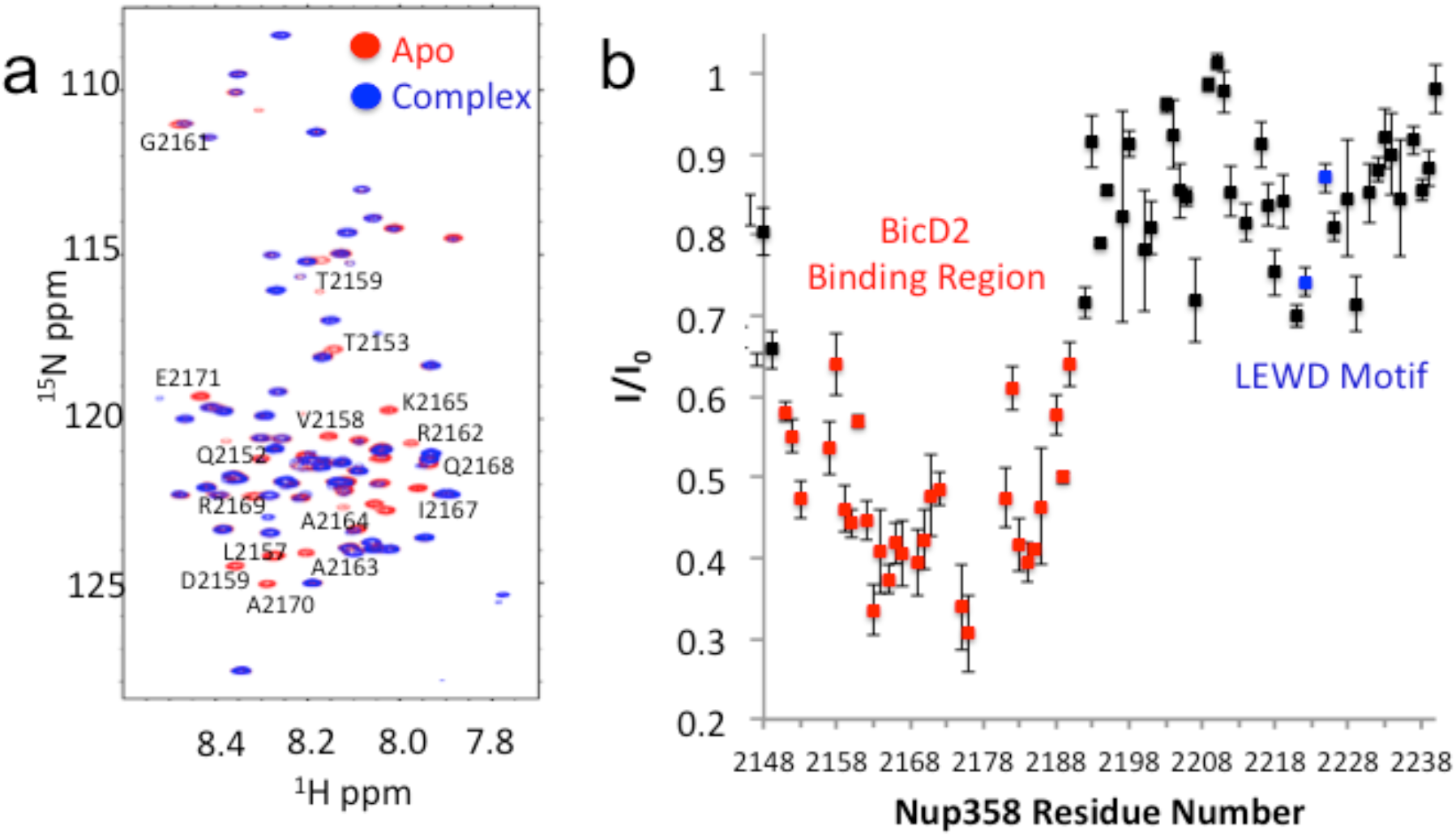
NMR titration narrowed the BicD2 binding site to the N-terminal half of Nup358-min. NMR mapping of Nup358 regions involved in BicD2-CTD binding was performed by titration of ^15^N labeled Nup358-min with BicD2-CTD. (a) The HSQC spectrum of a 1:1 Nup358-min/BicD2-CTD complex (blue) is overlaid on that of apo-Nup358-min (red). Many peaks disappeared in the complex spectrum (labeled by residue name and number), indicating that BicD2 binding causes slow to intermediate chemical exchange on the NMR time scale. (b) Plot of the peak intensity decrease vs the residue number of Nup358. The peak intensities of the 1:1 BicD2-CTD/Nup358-min complex spectrum were divided by the peak intensities of the apo-Nup358 spectrum to obtain I/I_0_. Data points with an I/I_0_ of 0.65 or lower are colored red. This plot shows that the N-terminal half of Nup358-min is largely responsible for BicD2 binding. The peak intensities corresponding to the LEWD sequence motif (colored blue) in Nup358, which mediates binding of KLC2, is not affected by BicD2 binding.

The Nup358-min domain also contains the previously published KLC2 binding LEWD motif^47^, but the BicD2 binding site determined here is separated from the LEWD motif by over 30 residues (Fig. 3b). Residues 2192 to 2240 of Nup358 show little change in NMR signal between the apo sample and the complex (Fig. 3b), indicating that the LEWD motif is not involved in BicD2 binding. Therefore, kinesin-1 and the dynein adaptor BicD2 may bind to separate but spatially close binding sites on the cargo adaptor Nup358. The close proximity of these motor recognition sites may serve to enable the interaction between dynein and kinesin machineries, for the regulation of bi-directional motility and precise positioning of the nucleus.

### CEST revealed a coil-to-helix transition at the BicD2-Nup358 interface

Although our results from the HSQC titrations establish the general region of the BicD2 interface in Nup358, the chemical shifts of the bound state were not obtained, due to peak disappearance in the HSQC upon BicD2 addition. To further characterize the BicD2/Nup358 interface with NMR, chemical exchange saturation transfer (CEST) experiments were performed. CEST has recently emerged as a powerful technique in solution NMR for measuring the chemical shifts of NMR invisible states^55,56^, such as the BicD2-CTD bound state of Nup358-min, whose resonances are absent from the ^15^N-HSQC (Fig. 3a, b). In CEST, when the invisible state is saturated by a weak and long radiofrequency (RF) pulse (B_1_), the saturation of the bound state will be transferred to the free state due to Nup358-min dissociating from the Nup358-min-BicD2 complex, causing a dip in peak intensity when B_1_ is on resonance with the bound state chemical shift, generating the minor dip in the CEST curve. If the chemical shifts of bound and unbound state are significantly different, which is for example the case for interface residues or residues that undergo structural changes upon complex formation, this difference gives rise to a double-dip appearance in the CEST profile (Fig. 4c and 4d). The major dip in the profile will be observed at the chemical shift of the free or unbound state (when the B1 matches the chemical shift of unbound resonance, the “visible state”). The second, smaller dip in the profile peak will be observed at the chemical shift of the bound state (due to saturation transfer from the “invisible state”). In contrast, residues that are not located at the complex interface or do not undergo structural changes upon complex formation will result in resonances without chemical exchange and the CEST profile will only show a single dip (Fig. 4a and 4b), corresponding to the chemical shift of the unbound state.

**Fig. 4.**
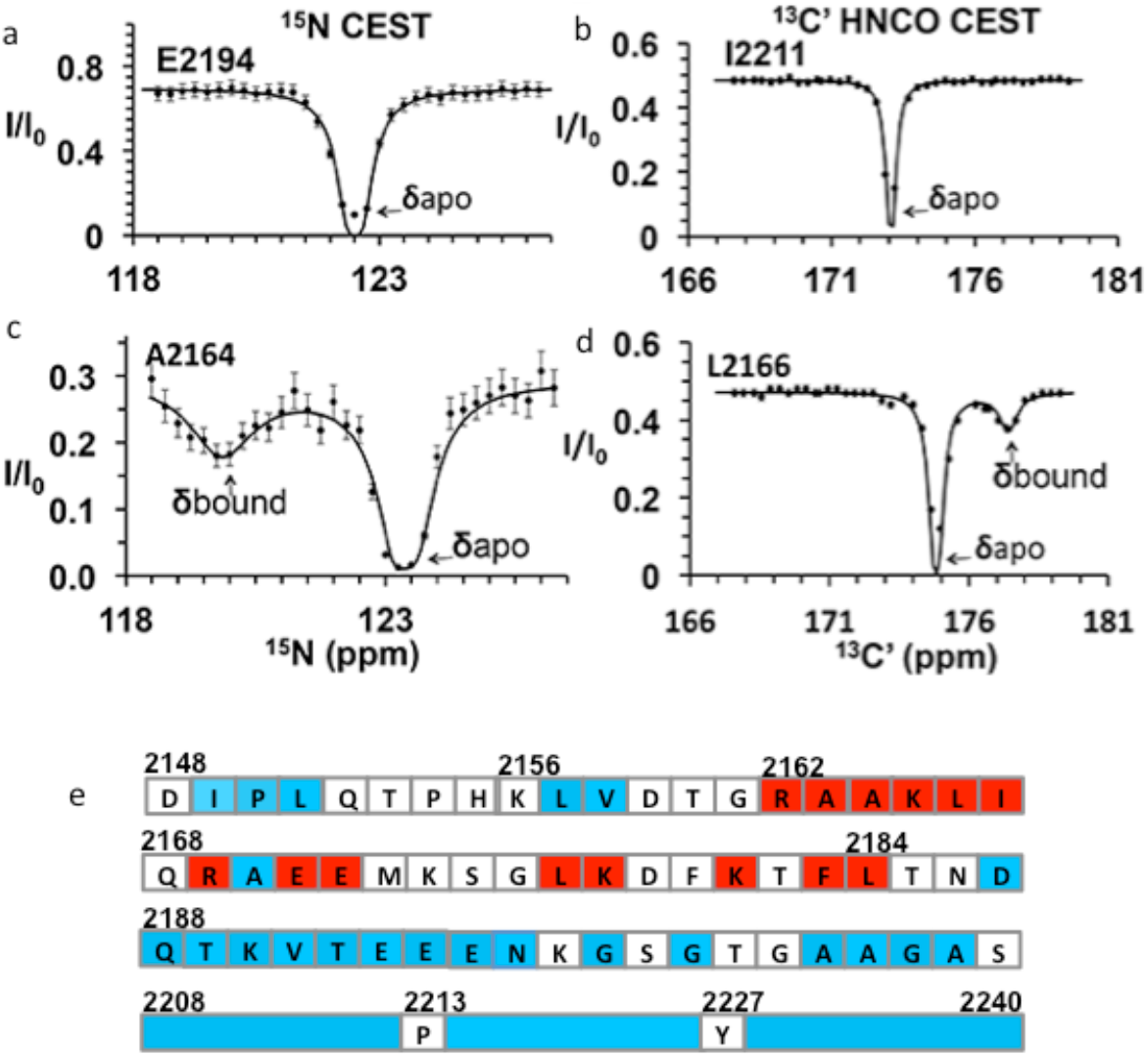
CEST maps chemical shifts of NMR-invisible, BicD2-bound state of Nup358. In the CEST profile curve, E2194 and I2211 have only a single dip in ^15^N-CEST (a) and ^13^C’-CEST (b), respectively, due to little chemical shift perturbation upon BicD2-CTD binding. This suggests E2194 is not at the binding interface and does not undergo conformational transition upon BicD2-CTD binding. In contrast, A2164 shows not only a dip at the chemical shift of the apo HSQC assignment, but also a smaller dip at about 3 ppm to the left in ^15^N-CEST (c), indicating that A2164 is in the binding region and/or experiences a conformational transition upon complex formation. (d) L2166 has a major peak and then a minor peak about 2.5 ppm to the right. The magnitude and direction of the chemical shift difference suggests a random coil to α-helical transition (see Table 1). (e) Summary of the CEST data obtained across the entire Nup358-min region. Red squares show residues with double dip in CEST profile. Their chemical shift changes from the apo to the bound state indicate that these red residues are part of the α-helical region at the Nup358/BicD2 interface. Blue colored residues show a single dip in CEST profile, suggesting these still remain in random coil conformation. White represents residues where data were not obtained, due to low signal to noise ratio or resonance overlap. All the curves with double dips and a sampling of curves with only single dip fits are shown in Fig. S3 through S6.

We performed CEST experiments on both the ^15^N amide and the ^13^C carbonyl (C’) resonances. In ^15^N CEST NMR experiments, the following residues were identified as interface residues or residues that undergo structural transition because of the double dip appearance in the CEST profile: A2163, A2164, K2165, L2166, I2167, K2178, L2184 (Fig. S3). For these residues, CEST indicates a significant chemical shift difference between their bound and unbound states. Using the program RING NMR Dynamics^57^, the CEST data were fit to obtain chemical shift difference between free and bound state (Δδ), the exchange rate (k_ex_) and the fractional minor population (P_b_)(See Table S1a, b). The weighted average of the ^15^N CEST fit parameters resulted in k_ex_ = 200 ± 70 s^−1^, consistent with slow to intermediate exchange (k_ex_ < Δω) on an NMR time scale. The weighted average of the minor population (P_b_) is 0.04 ± 0.01, which is consistent with our sample preparation at molar ratio of 20:1 for BicD2-CTD: ^15^N-Nup358^min^. Many peaks in the ^15^N-CEST suffered from low-signal-to-noise ratio and resonance overlap common in IDPs, preventing the determination of additional chemical shifts of invisible states. To overcome this problem, we further carried out ^13^C’-CEST experiments based on HNCO, which correlates amides (HN) with the carbonyl (CO) of the ***preceding*** residue. Here the saturation pulse B_1_ is on the carbonyl, and intensity change due to saturation is still reported in an ^15^N-HSQC-like 2D spectrum. This method provided additional data for residues missed with the ^15^N CEST due to resonance overlap or low signal to noise ratio. For example, the F2183 peak in ^15^N-^1^H HSQC was overlapped. However, the L2184 peak, a well-resolved signal with high S/N, showed a CEST effect from the saturation of the F2183 carbonyl in HNCO-based ^13^C’-CEST. With the ^13^C HNCO-CEST experiment, we were able to observe minor states in the following residues – R2162, A2164, K2165, L2166, R2169, E2171, E2172, L2177, K2181, F2183, and L2184 (Fig. S4 and Table S1c). Thus, with CEST, we were able to further narrow the binding region to residues 2162 through 2184 (Fig. 4e). Fitting of HNCO-CEST profiles with RING Dynamics resulted in a global exchange rate of 270 ± 50 s^−1^, in reasonable agreement with the value of k_ex_ from ^15^N CEST. The weighted average of the minor population (P_b_) is 0.06 ± 0.01, which is again consistent with our sample preparation. The 18 CEST profiles with double dip appearance all resulted in chemical shift changes in the direction and magnitude as expected for a random coil to α-helix transition^58^ (typically the chemical shift for ^15^N becomes lower in the helical conformation, and higher for ^13^C, as shown in Table 1). The CEST results showed that Nup358 residue 2162-2184 undergoes coil to helix transition upon binding to BicD2, suggesting that BicD2 recognizes Nup358 through a short “cargo recognition α-helix” which is embedded in an IDP domain.

**Table 1:**
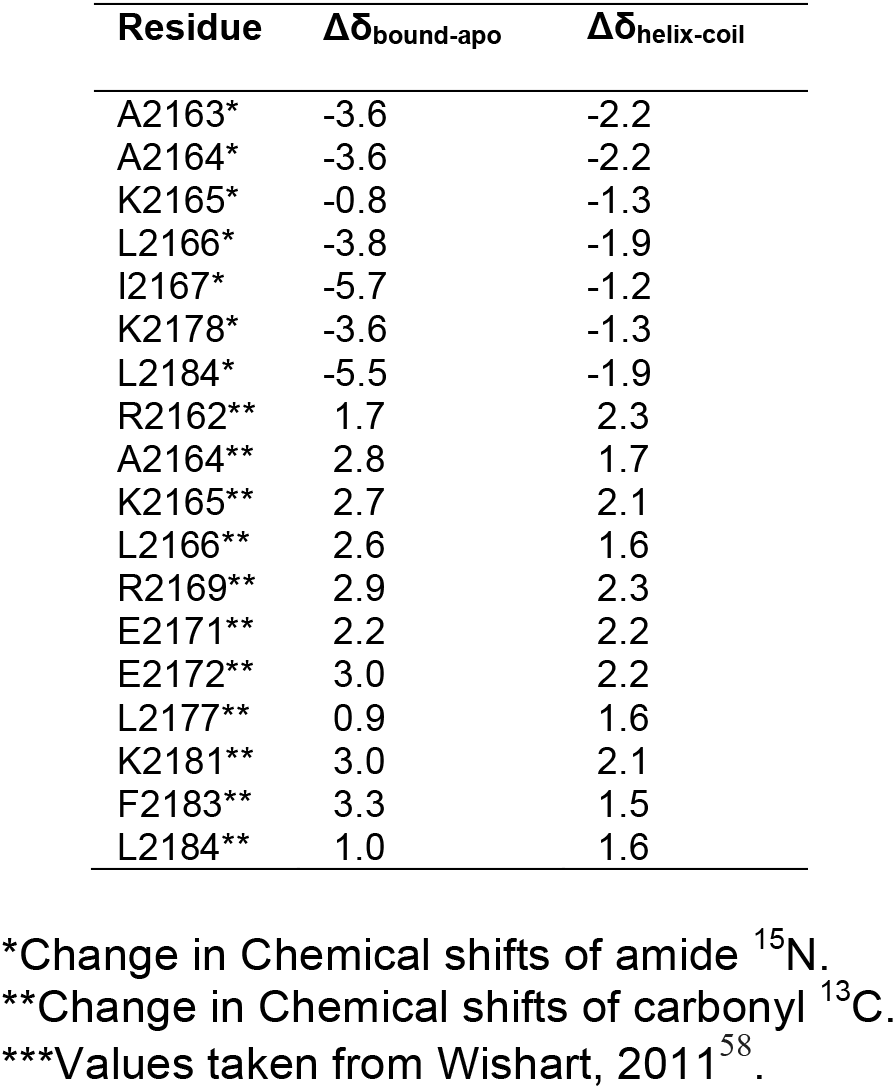
Chemical shift differences from CEST of Nup358/BicD2 and apo-Nup358 (Δδ_bound-apo_) match closely to Δδ for coil to helix transition (Δδ_helix-coil_)***.

| Residue | $\Delta\delta_{\text{bound-apo}}$ | $\Delta\delta_{\text{helix-coil}}$ |
| --- | --- | --- |
| A2163* | -3.6 | -2.2 |
| A2164* | -3.6 | -2.2 |
| K2165* | -0.8 | -1.3 |
| L2166* | -3.8 | -1.9 |
| I2167* | -5.7 | -1.2 |
| K2178* | -3.6 | -1.3 |
| L2184* | -5.5 | -1.9 |
| R2162** | 1.7 | 2.3 |
| A2164** | 2.8 | 1.7 |
| K2165** | 2.7 | 2.1 |
| L2166** | 2.6 | 1.6 |
| R2169** | 2.9 | 2.3 |
| E2171** | 2.2 | 2.2 |
| E2172** | 3.0 | 2.2 |
| L2177** | 0.9 | 1.6 |
| K2181** | 3.0 | 2.1 |
| F2183** | 3.3 | 1.5 |
| L2184** | 1.0 | 1.6 |

\*Change in Chemical shifts of amide $^{15}\text{N}$ .
\*\*Change in Chemical shifts of carbonyl $^{13}\text{C}$ .
\*\*\*Values taken from Wishart, 2011<sup>58</sup>.

### CD spectroscopy confirms formation of anα-helix in Nup358 upon binding to BicD2

To provide further evidence for the coil-to-α-helix transition,circular dichroism (CD) spectroscopy was carried out. The CD wavelength scans of the Nup358-min/BicD2-CTD complex had minima at 208 nm and 222 nm, characteristic for α-helical proteins ^30^ (Fig. 5). Such minima were absent in the CD spectra of Nup358-min, as expected for an intrinsically disordered protein. Notably, the minima at 208 and 222 nm were shifted towards more negative values in the complex compared to the sum of the individual spectra of Nup358-min and BicD2-CTD, suggesting that the α-helical content increases upon complex formation.

**Fig. 5.**
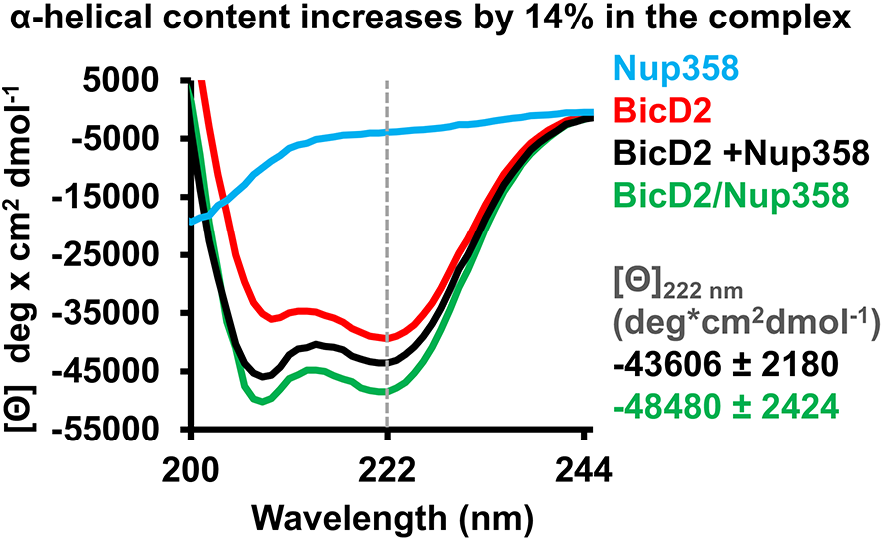
CD spectroscopy confirms formation of an α-helix in the Nup358/BicD2 complex. CD wavelength scans of BicD2-CTD (red), Nup358-min (blue) and the Nup358-min/BicD2 complex (green) at 4°C are shown. The sum of the individual wavelength scans of Nup358-min and BicD2-CTD is shown in black. The mean residue molar ellipticity [Θ] versus the wavelength is shown. Experiments were repeated two or three times, representative scans are shown. A characteristic feature of α-helical structures is a local minimum at 222 nm (dashed line). Based on the values for [Θ] at 222 nm (shown on the right) and based on our calibration curve (Fig. S7), the Nup358/BicD2 complex has a 14 ±5 % increase in α-helical content compared to the sum of the spectra of BicD2 and Nup358.

To quantify the increase of the α-helical content, we determined the difference between the molar ellipticity at 222 nm of the complex and of the sum of the individual proteins, Δ[Θ], to be −4874 deg*cm^2^ dmol^−1^ (Fig. 5), which corresponds to a 14 ±5 % increase of the α-helical content based on a published thermal unfolding curve of the BicD2-CTD, which was used for calibration^30^ (Fig. 5, Fig. S7 and ref. ^59^). We recently determined the experimental error to be ~5% (corresponding to Δ[Θ] of 2930 deg*cm^2^ dmol^−1^)^59^; therefore, we estimate that the α-helical content increases by 14 ± 5 % upon Nup358/BicD2 complex formation, confirming a structural transition from random coil to α-helix.

### The minimal Nup358/BicD2 complex has a rod-like shape that is more compact than individual proteins

Small-Angle X-ray Scattering (SAXS) experiments were carried out to obtain a low-resolution structure of the Nup358-min/BicD2-CTD complex. The quality of our SAXS data was confirmed by molar mass calculations: for Nup358-min, we determined a molar mass of 12.3 kDa (Table S2), which matches closely to the calculated mass of a monomer (10.6 kDa). For the Nup358-min/BicD2-CTD complex, we determined a molar mass of 47.6 kDa (Table S2), which matches closely to the expected mass of a Nup358-min/BicD2-CTD complex with a 2:2 stoichiometry (calculated molar mass of 43.0 kDa). This is in line with our previously published SEC-MALS data which suggest that apo Nup358-min forms monomers, while the Nup358/BicD2 complex has a 2:2 stoichiometry^47,50^.

A Kratky plot of the SAXS profiles further confirmed that Nup358-min is intrinsically disordered, as the signal increases at high q values instead of approaching zero (Fig. 6a). In contrast, the Kratky plots of the BicD2-CTD and the Nup358-min/BicD2-CTD are bell-shaped, suggesting they are folded (Fig. 6a). Thus, Nup358-min becomes more compact upon complex formation with BicD2-CTD, consistent with the coil-to-helix transition observed from NMR and CD.

**Fig. 6.**
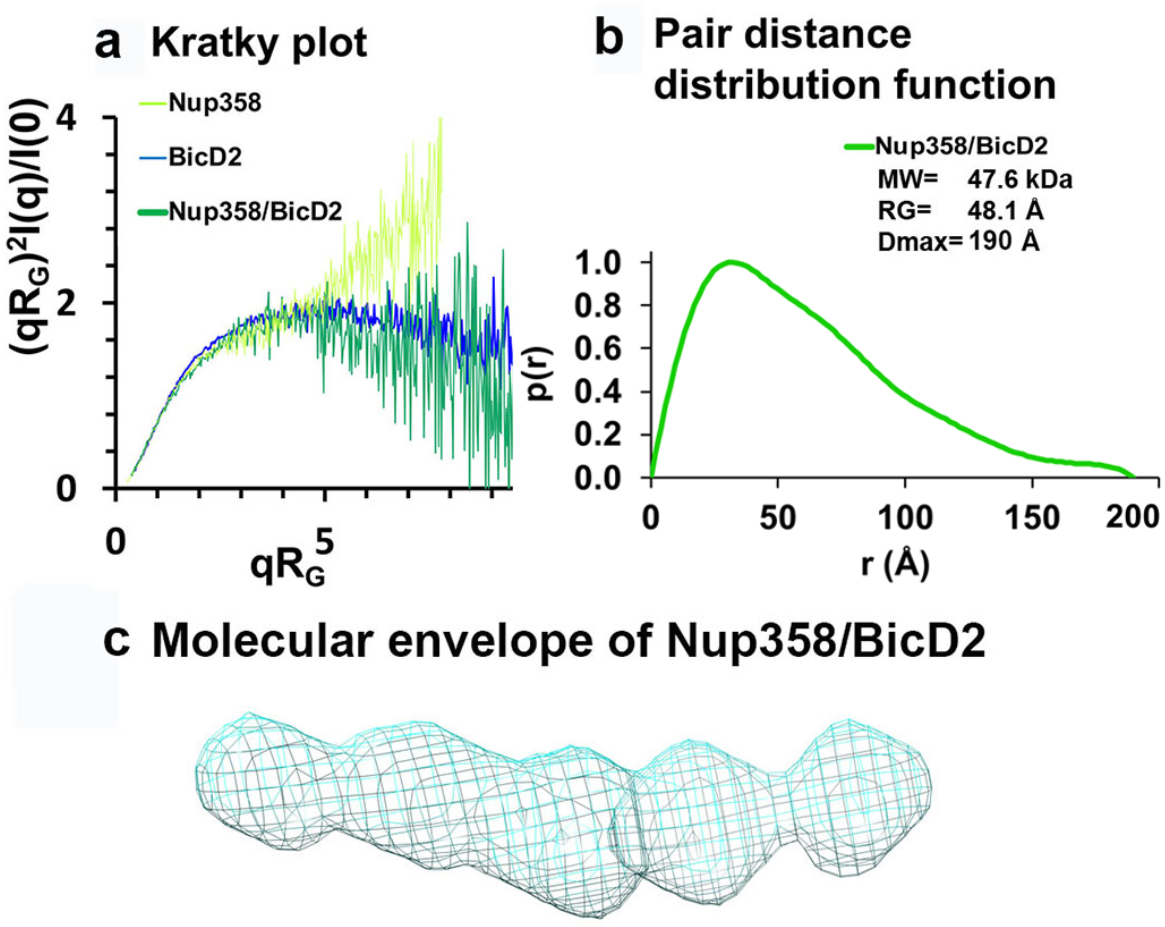
Low resolution structures determined by SAXS confirm that the complex has a rod-like shape that is more compact than the individual proteins. (a) Dimensionless Kratky plots of the SAXS data collected from the minimal Nup358/BicD2 complex, from Nup358-min and BicD2-CTD (q: scattering vector, R_G_: radius of gyration, I(q): scattering intensity). (b) The pair distance distribution function, p(r), of the Nup358-min/BicD2-CTD complex was derived from the scattering intensity profile (Fig. S8). The molecular weight MW^60^, the radius of gyration R_G_ from the Guinier plot and the largest dimension of the particle D_max_ are shown. (c) Refined bead model 3D reconstruction of the Nup358-min/BicD2-CTD complex (cyan mesh). The statistical analysis and supporting SAXS data are summarized in Table S2 and Figs. S8-11.

The pair distance distribution function (P(r)) derived from the SAXS profile of the Nup358-min/BicD2-CTD complex has a peak which decays with a linear slope, which is characteristic for elongated, rod-like structures (Fig. 6b). Since the width of a rod remains similar throughout the rod, the sum of all pair distances will have a linear slope. The maximum particle diameter D_Max_ of the complex is 190 Å, which is identical to the D_Max_ that was determined for the BicD2-CTD sample (Table S2), suggesting that the overall length of the rod-structure does not change. Bead models of the Nup358-min/BicD2 complex were reconstructed from the P(r) functions and confirm that the complex has a flexible, rod-like structure (Fig. 6c). The normalized spatial discrepancy (NSD) is 0.7, suggesting the structural convergence and homogeneity of bead models reconstructed from SAXS profiles (Table S2).

Taken together, these data suggest that the complex has a 2:2 stoichiometry and is more compact than the apo state, further validating the coil-to-helix transition in the Nup358/BicD2 complex.

### Nup358 mutagenesis validates α-helix formation at the Nup358/BicD2 interface

Our CEST, CD and SAXS data provided strong evidence that residues 2162-2184 of Nup358 undergo a structural transition from random coil to α-helix upon complex formation with BicD2. In addition, residues 2162-2184 of Nup358 are also predicted to form an α-helix by secondary structure prediction^61^.

To confirm our results, we also carried out alanine mutagenesis for all residues of the Nup358 α-helix and assessed binding of the mutated GST-tagged Nup358-min to BicD2-CTD by pulldown assays (Fig. 7 and Fig. S12). Eleven mutants displayed diminished binding, which are interspaced throughout the α-helix. An α-helical wheel representation was created, which revealed that these interface residues are clustered on one side of the α-helical wheel (red in Fig. 7c). The mutagenesis experiments are consistent with our results from NMR spectroscopy. While interfacial residues and residues that undergo the transition from random coil to the α-helix will show a chemical shift change, not all residues in the new α-helix in Nup358-min will be at the binding interface, therefore the mutagenesis data provides additional information. Notably, all interface residues that were identified from mutagenesis (red in Fig. 7d) also had a double dip in the CEST profile (red in Fig. 7d) or were not assessed (white in Fig. 7d). Furthermore, removal of residues 2163-2166 (AAKL) of Nup358-min (ΔAAKL), which had double dips in CEST curves, virtually abolishes the interaction with the BicD2-CTD (Fig. S12b). We also removed the α-helix from Nup358-min, and as expected, the resulting Nup358-fragment (residues 2185-2240) shows virtually no interaction with the BicD2-CTD (Fig. S12 a, last lane). Finally, we successfully assembled a minimal complex of the BicD2-CTD with a Nup358 fragment (residues 2157-2199) that only contained the BicD2 binding site from NMR, which confirmed our mapped binding site (Fig. S12c). Furthermore, known BicD2 residues that are important for Nup358 interaction include L746, R747, M749 and R753^29^, suggesting a potential binding site for the Nup358 α-helix on the BicD2-CTD coiled coil.

**Fig. 7.**
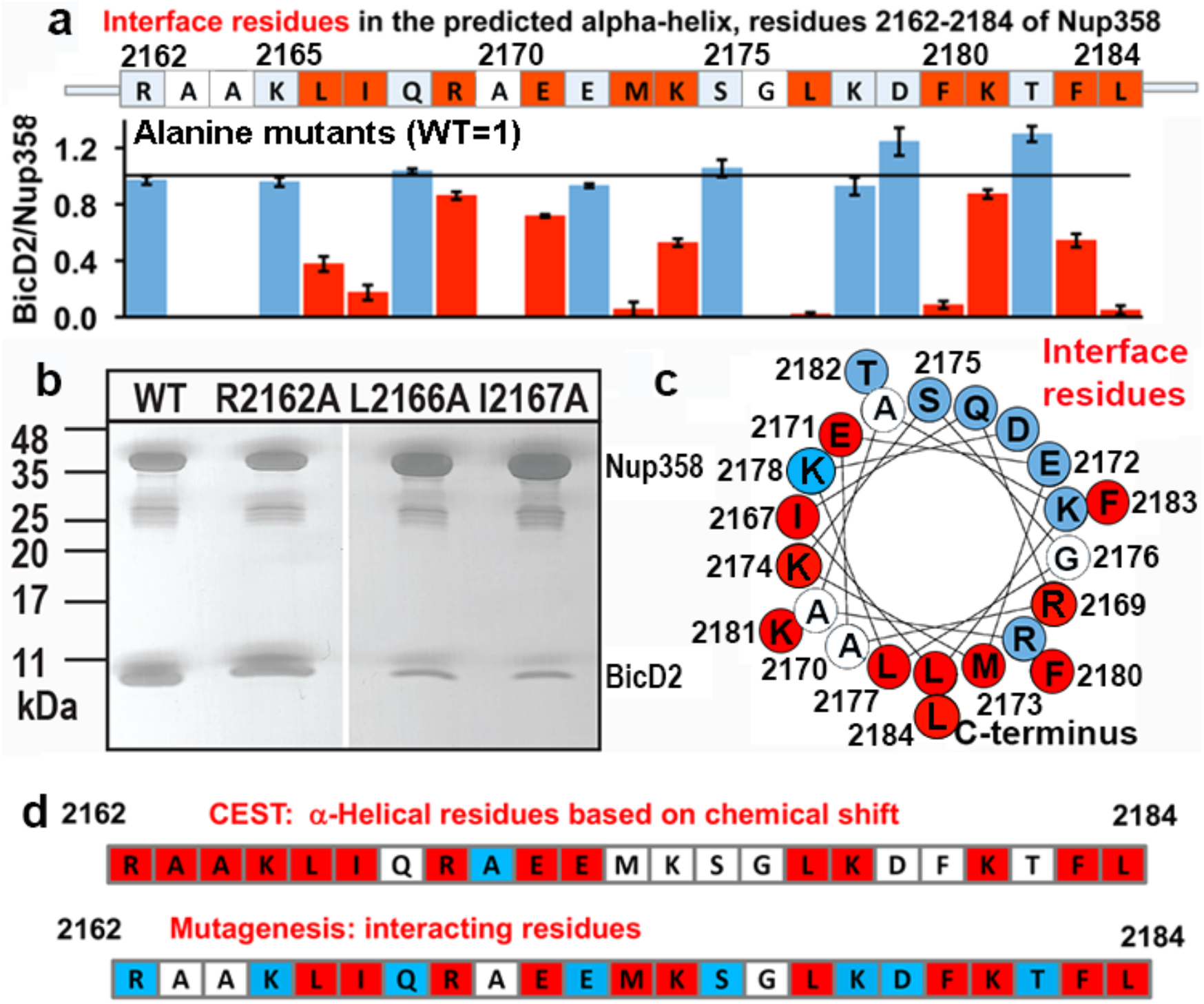
Mutagenesis of the Nup358 cargo recognition helix. All residues of the Nup358 α-helix were mutated to alanine and binding to BicD2-CTD was assessed by pull-down assays. Nup358-min-GST was immobilized on glutathione sepharose beads and incubated with purified BicD2-CTD prior to elution with glutathione. The elution fractions were analyzed by SDS-PAGE and the intensities of the gel bands were quantified to obtain the ratio of bound BicD2-CTD/Nup358-min, normalized respective to the wild-type (Fig. S12). Eleven interface residues were identified, which are colored red and show a reduction of binding upon mutation by at least three times the standard deviation. Alanine and glycine residues were not assessed and are colored white. Residues for which mutations do not affect binding are colored light blue. (a) The sequence of the predicted α–helix is shown above a bar graph of the ratios of bound BicD2 to Nup358 from the alanine mutant pull-down assays, normalized respectively to the WT (WT=1, indicated by the horizontal black line). The ratios were averaged from three independent experiments and the standard deviations were calculated as errors. (b) Representative SDS-PAGEs of elution fractions of the pull-down assays. The gel band intensities were quantified. A representative full dataset is shown in Fig. S12. Note that a small gel band at 25 kDa represents GST. (c) Helical wheel representation for (a). (d) Comparison of helical residues in Nup358-BicD2 complex identified by CEST and results from pulldown assay of mutants. Red: CEST positive (double dip in CEST profile) or strong reduction in binding from the pulldown assay with alanine mutation at this residue; blue: CEST negative (single dip in CEST profile) or no effect in mutagenesis; white: data not available.

### Nup358-BicD2 interface is essential for dynein activation

Single-molecule binding and processivity assays were used to further explore the impact of Nup358 mutants on dynein recruitment and activation of dynein motility. We assessed the interaction between BicD2 and the following Nup358^min-zip^ mutants: I2167A, M2173A, L2177A, F2180A, and L2184A, in the context of the BN^min-zip^ and the DDBN^min-zip^ complex. The following parameters were quantified for WT and mutant Nup358^min-zip^ constructs: (a) binding frequency of DDBN^min-zip^ to MTs in the absence of MgATP, (b) percent complex formation with BicD2 alone or to the DDB complex, and (c) speed and run length of DDBN^min-zip^ on MTs (Fig. 8, Fig. S14).

**Fig. 8.**
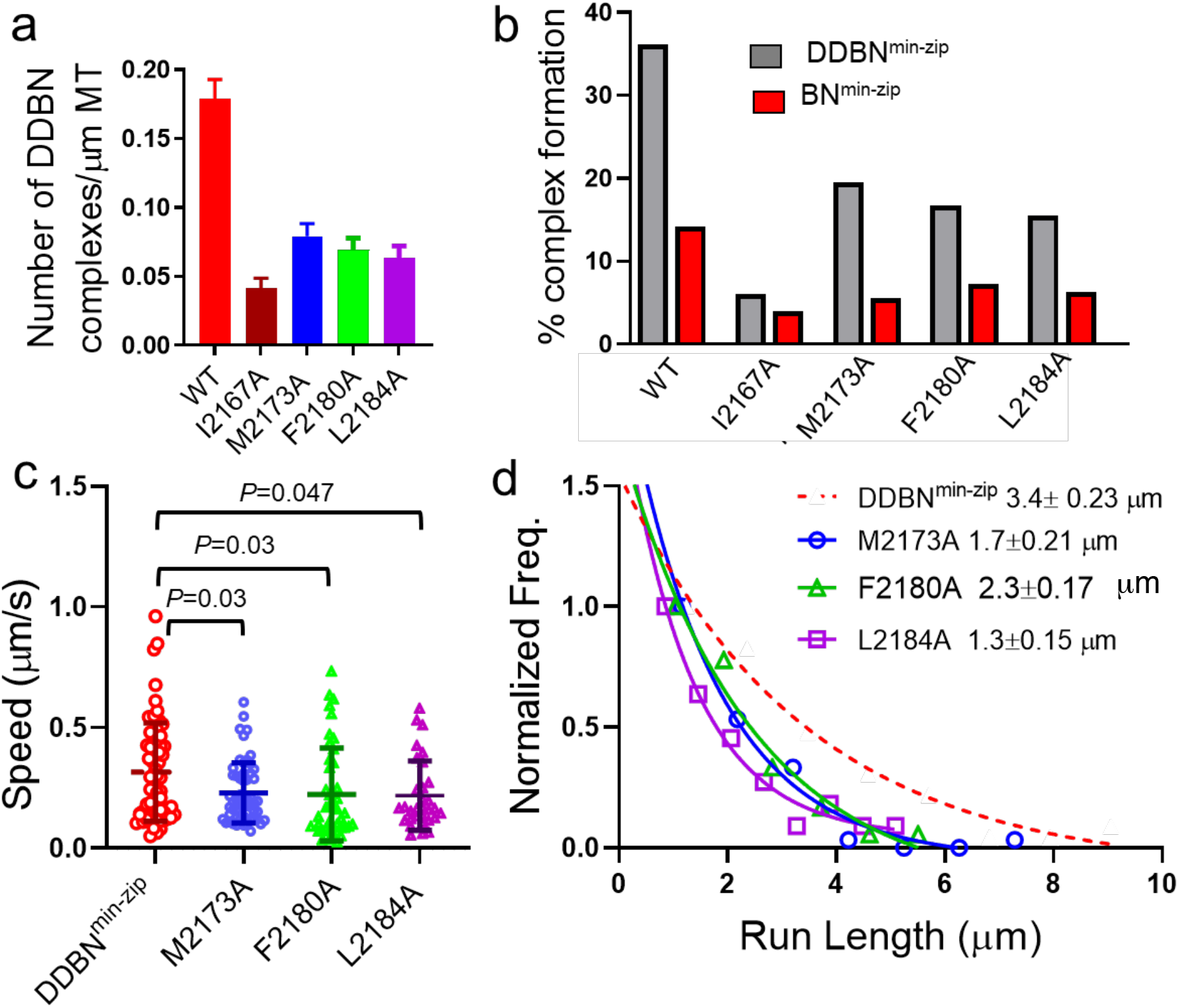
Nup358 point mutations that diminish the interaction with BicD2-CTD also diminish formation of the DDBN complex. (a) Histogram of the binding frequency of DDBN^min-zip^ complexes per micrometer MT length. The binding frequency of DDBNmin-zip complexes formed with WT-Nup358^min-zip^ is significantly higher (p<0.001) than those formed with the Nup358^min-zip^ mutants I2167A, M2173A, F2180A, and L2184A. The binding frequency of DDBN^min-zip^ formed with WT-Nup358^min-zip^ was 0.18±0.014/μm (n=29, with n being the number of microtubules). In contrast, these values for DDBN^min-zip^ formed with the Nup358^min-zip^ mutants I2167A, M2173A, F2180A, and L2184A were 0.041±0.007/μm (n=29), 0.079±0.009/μm (n=30), 0.069±0.009/μm (n=28), 0.063±0.008/μm (n=28) respectively. b) The presence of dynein-dynactin (gray) increases the formation of BicD2-Nup358^min-zip^ complexes (red). The percent formation of BN^min-zip^ was: 14.13% (n=580) for WT, 3.94% (n=1142) for I2167A, 5.54% (n=1877) for M2173A, 6.3%(n=412) for F2180A and 7.2% (n=1032) for L2184A Nup358^min-zip^. In contrast, these values for DDBN^min-zip^ complexes were 36.1% (n=945), 6.09% (n=986), 19.5% (n=347), 15.5%(n=890) and 16.66% (n=372), respectively. (c) Comparison of the speeds of DDBN complexes formed with varying Nup358^min-zip^ constructs. The speed of DDBN formed with Nup358^min-zip^ mutants M2173A (0.23 ±0.12 μm, N=58; blue), F2180A (0.22±0.19 μm, n=43; green) and L2184A (0.21±0.14 μm, n=31; pink) are significantly slower than that of DDBN-WT (0.32±0.20, n=68; taken from Fig. 2h) (*p*= 0.03, 0.03 and 0.047 respectively; one-way ANOVA followed by Tukey’s test). (d). The run length of DDBN^min-zip^ formed with Nup358^min-zip^-M2173A (1.7±0.21 μm/s, n=58), F2180A (2.3±0.17 μm/s, n=43) and L2184A (1.3±0.15 μm/s, n=31) are significantly shorter than that formed with WT-Nup358^min-zip^ (3.4±0.23 μm/s; n=68; taken from 2i) (with *p-values* p=0.009, 0.04, and p=0.048 respectively; one-way ANOVA followed by Tukey’s test).

The binding frequency of DDBN^min-zip^ complexes to the MT was calculated by counting the number of dual-color Qdots bound per MT length (Fig. 8a). The binding frequency of DDBN^min-zip^ formed with the Nup358^min-zip^ mutants I2167A, M2173A, F2180A, and L2184A Nup358^min-zip^ was significantly lower than that formed with WT-Nup358^min-zip^ (Fig. 8a).

We next assessed how these four Nup358^min-zip^ mutants affected the BicD2-Nup358^min-zip^ (BN) interaction in the absence of dynein-dynactin. BicD2 and WT or mutant Nup358^min-zip^ were labeled with different color Qdots and the number of dual-color complexes on the glass surface quantified using TIRF microscopy (Fig. S14) (see Methods). Nup358^min-zip^ binds BicD2 alone poorly, and the percentage of BN was reduced further for four Nup358^min-zip^ mutants (Fig. 8b). We also quantified how the presence of dynein-dynactin influences the BicD2-Nup358^min-zip^ interaction. Strikingly, in the presence of dynein-dynactin, the formation of DDBN^min-zip^ complexes was about 2-3 times higher than the BN complexes (Fig. 8); but again the four mutant Nup358^min-zip^ showed reduced complex formation relative to WT (Fig. 8b). The difference between DDBN^min-zip^ and BN^min-zip^ indicates that dynein-dynactin enhances the BicD2-Nup358 interaction.

The motion of DDBN^min-zip^ complexes formed with these four Nup358^min-zip^ mutants was quantified. Complexes with the Nup358^min-zip^ mutants I2167A, M2173A, F2180A and L2184A all showed slower speed and shorter run-length compared with the WT-DDBN^min-zip^. Data obtained with the DDBN^min-zip^ complex containing either WT, M2173A, F2180 A or L2184A Nup358^min-zip^ are shown (Fig. 8c, d). There were too few dual-colored complexes formed with the I1267A mutant to allow speed and run length quantification. Compromised motility in the presence of Nup358^min-zip^ mutants suggests formation of a less stable DDBN complex.

Although four Nup358^min-zip^ complexes showed significantly reduced MT binding affinity, speed and run length compared to the WT-Nup358^min-zip^, unexpectedly, DDBN^min-zip^ complex formed with the mutant Nup358^min-zip^-L2177A behaves similarly to WT-Nup358^min-zip^ (Fig. S13). The MT binding frequency, speed and run length of DDBN^min-zip^_complex formed with Nup358^min-zip^-L2177A was the same as that formed with WT-Nup358^min-zip^ (Fig. S13). The results with the Nup358^min-zip^-L2177A mutant imply that there is some plasticity at the Nup358/BicD2 interface. It is striking that four out of the five point mutants tested from the cargo recognition helix significantly impaired DDBN^min-zip^ complex formation and processive motility, validating the functional importance of this helix.

## Discussion

Dynein mediated nuclear positioning plays crucial roles in many fundamental biological processes, such as mitosis, meiosis, brain and muscle development^2–16,62^. However, the molecular mechanisms for cargo recognition by dynein adaptor proteins remain poorly understood. Here, we used multiple biophysical and biochemical methods to elucidate the molecular details of the interface between dynein adaptor BicD2 and the cargo adaptor Nup358. First, we identified a minimal complex between BicD2-CTD and Nup358-min, with μM affinity. Single-molecule processivity assays revealed that a dimerized Nup358-min domain forms a complex with dynein/dynactin/BicD2 (DDBN) that is activated for processive motility on microtubules. The observation that dimerization of Nup358-min enhances the interaction with BicD2 and with the DDB complex suggests that the activation is linked to oligomerization. We next proceeded to characterize the structure of the minimal complex with NMR, CD and SAXS. NMR titration of BicD2-CTD into ^15^N labeled Nup358-min narrowed the binding region to be in the N-terminal half of Nup358-min. However, due to slow chemical exchange on the NMR time scale and fast relaxation in the bound state, the BicD2-bound state of Nup358-min cannot be directly observed in NMR, as in an NMR “invisible state”, or “dark state”. Therefore, a powerful solution NMR technique, CEST, was applied to map the chemical shifts of the “invisible state”. The CEST-derived bound state chemical shifts of Nup358-min not only identified key residues at the Nup358/BicD2 interface, but also showed that chemical shift changes upon binding correspond to that of a coil-to-helix transition in Nup358-min. This conformation change was confirmed by CD spectroscopy; furthermore, SAXS experiments confirmed that the Nup358-min/BicD2-CTD complex has a 2:2 stoichometry^50^ and is more compact than the individual proteins. The α-helical interface was also validated by mutagenesis, which identified interface residues that form a stripe along the helical surface. Single molecule binding assays with five of these mutants confirmed that they all diminish the interaction of Nup358-min with full-length BicD2. Strikingly, in four out of the five point mutations in Nup358^min-zip^ also decrease dynein/dynactin/BicD2/Nup358 (DDBN) complex formation, and the complexes that were formed exhibited reduced run length and speed reflecting formation of a less stable complex. Our data highlight the important role of dynein adapter/cargo adapter interactions in activating and fine-tuning dynein motility. We also observed an exception in L2177A which behaved most similarly to WT in single molecule experiments, suggesting some plasticity at the Nup358/BicD2 interface.

Our data establish that BicD2 recognizes its cargo Nup358 though an α-helix of ~28 residues, which may be a structural motif that stabilizes BicD2 in its activated state. Coil-to-helix transition and folding upon binding, as shown here, have been observed in many studies of IDP/IDR interactions and recognition^40,56,63^. Our single molecule motility studies of the WT and mutant Nup358^min^ further confirm the significance of the Nup358 cargo recognition α-helix for dynein recruitment and motility. We therefore propose that *the dynein adaptor BicD2 recognizes its cargos through a “cargo recognition α-helix”* (Fig. 9). It is conceivable that the cargo adaptors Rab6^GTP^ and nesprin-2G are also recognized by similar short α-helices, although this needs to be confirmed by future experiments. Our data thus suggest a structural basis for cargo recognition by BicD2 and will enable further studies aimed at identifying cell cycle specific regulatory mechanisms for distinct transport pathways that are facilitated by BicD2^4,11,50^.

**Fig. 9.**
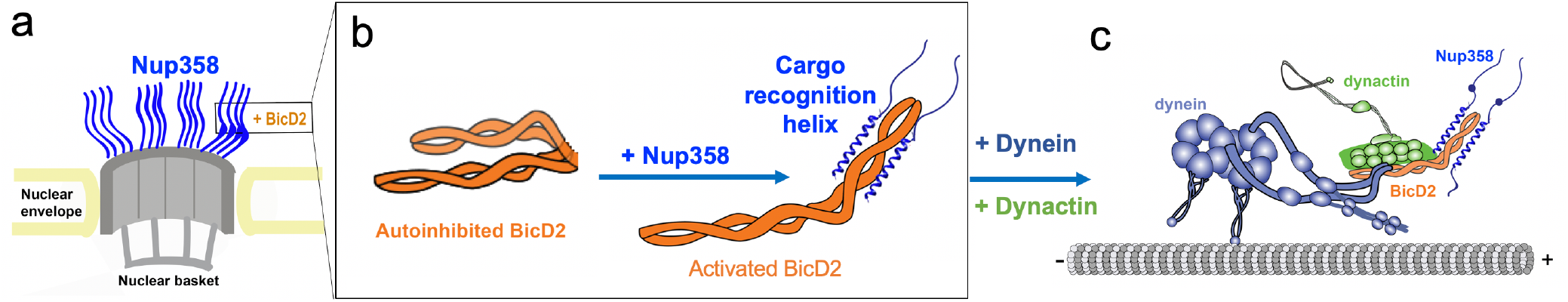
We propose that BicD2 recognizes its cargo through a short “cargo-recognition α-helix”, which may also be a structural feature that stabilizes the activated state of BicD2 for the recruitment of dynein and dynactin. (a) Cutaway view of half of an NPC. Each of the eight spokes of the NPC contains four molecules of Nup358 on the cytoplasmic side (i.e., 32 in total), which provide binding sites for dynein (via BicD2) and kinesin-1 (via KLC2). Nup358 makes up most of the mass of the cytoplasmic filaments of the NPC^77,78^.(b) Left panel: Schematic representation of the looped, auto-inhibited conformation of BicD2 which is formed in the absence of cargoes such as Nup358. Apo-Nup358 contains many intrinsically disordered regions (IDR), including the BicD2 binding region. Upon binding of Nup358 to BicD2, a short α-helix is formed at the Nup358/BicD2 interface, which is a structural feature that stabilizes BicD2 in the active state. Binding of Nup358 to BicD2 likely promotes loop opening, which activates BicD2 for dynein/dynactin recruitment^24^. (c) Schematic representation of the proposed dynein/dynactin/BicD2/Nup358 complex (DDBN) bound to microtubules for processive motility for nuclear positioning. Nup358 has a LEWD motif which recruits kinesin-1 via KLC2. The binding sites for the opposite polarity motor kinesin-1 on Nup358 are indicated by small blue sphere. The location of the LEWD motif which recruits kinesin-1 via KLC2 is indicated. We propose that BicD2 and KLC2 interact simultaneously with Nup358 for precise control of nuclear positioning.

A key step in activating dynein motility is the activation of BicD2 for dynein recruitment, and we and others have recently proposed that cargo-binding activates BicD2 for dynein recruitment by inducing a coiled-coil registry shift in BicD2 (i.e. a vertical displacement of the two α-helices against each other by one helical turn)^26,29,30,59^. In the absence of cargo, BicD2 forms an auto-inhibited looped dimer that cannot recruit dynein, with the CTD masking the N-terminal dynein/dynactin-binding site (NTD)^17–27,64^. When cargo is loaded, BicD2 can directly link dynein with its activator dynactin, which is required to activate dynein for processive motility^17–27,64^. Our recent work showed that an F684I mutant of the *Drosophila (Dm)* homolog BicD can activate dynein/dynactin for processive motility in the absence of cargo. X-ray structures of the WT and F684I mutant as well as molecular dynamics simulations suggest that this activating mutation causes a coiled-coil registry shift in the *Dm* BicD-CTD^30,59^. It is very tempting to hypothesize that the cargo recognition helix of Nup358, intercalates into the coiled-coil of the BicD2-CTD to stabilize a coiled-coil registry shift in the Nup358/BicD2 complex, resulting in activation of BicD2 for dynein recruitment. This hypothesis remains to be experimentally confirmed.

Interestingly, Nup358-min binds more strongly to the dynein/dynactin/BicD2 complex compared to full-length BicD2 alone (Fig. 8b). This observation supports the proposed loop-opening activation mechanism of BicD2^24–26,59^ (Fig. 9b, c). In the BN complex, the NTD of BicD2 competes with Nup358 for binding to the CTD, likely resulting in a lower apparent affinity than in the DDBN complex, where the NTD interacts with dynein and dynactin and prevents re-association of the NTD with the CTD. It is also possible that additional weak interfaces further stabilize the DDBN complex compared to Nup358/BicD2 or compared to the DDB^CC1^ complex. Here we observed a direct interaction of Nup358 with dynein-dynactin that could contribute to enhanced affinity. It has been shown for several cargo adaptors that they interact weakly with the dynein light chains^65^ and the dynein light intermediate chain 1 (LIC1) forms a small interface with the NTD of BicD2, which is required for activation of processive motility^40,41,63^. Cargoes and adaptors often interact through multiple interfaces with motors, and these additional interfaces in many cases enhance overall motility^2,22,23,40,41,43,66,67^.

Furthermore, the affinity of the monomeric cargo adaptor Nup358-min towards BicD2 is enhanced when it is dimerized by a leucine zipper, similar to what was previously observed for binding of the cargo adaptor Egalitarian for *Drosophila* BicD^24,25^. Thus, it is conceivable that a Nup358-min dimer binds to a BicD2-CTD dimer, which is also in line with our ITC results, which reveal a single K_D_. While Nup358-min is a monomer on its own, it oligomerizes to form 2:2 complexes with either the BicD2-CTD or the KLC2, and also forms a ternary complex with both BicD2-CTD and KLC2 with 2:2:2 stoichiometry ^47,50^. We thus propose that cargo adaptor-induced dimerization may potentially be a universal feature of the activation mechanism, since BicD2 and KLC2 both form dimers in active dynein and kinesin-1 motors^21,22,47,68^. Importantly, NMR titration showed that BicD2 binding has no effect on the NMR signal of the LEWD motif of Nup358 (Fig. 3), which can interact directly with KLC2^47^. These data further suggest that the Nup358-min domain is potentially capable of recruiting both dynein and kinesin-1 machineries for bi-directional positioning of the nucleus, although this remains to be confirmed in the context of full-length proteins and intact motors. Intriguingly, our NMR data show that the BicD2 binding site is only separated by 30 residues from the LEWD motif that acts as KLC2 binding site. The close proximity of these binding sites may play a role in coordinating motility for bi-directional transport^2–4,6,9,10,15,47,69–76^. Another intriguing possibility is that if kinesin binding dimerizes Nup358, this may be the key initial step leading to BicD2 activation and recruitment of dynein-dynactin to form a bidirectional complex (Fig. 9). The ternary Nup358/BicD2/KLC2 complex may be a model for other transport modules, in which opposite polarity motors dynein and kinesin-1 act together.

## Conclusion

Based on our data, we propose a structural basis for cargo recognition through the dynein adaptor BicD2. Our results establish that BicD2 recognizes its cargo Nup358 through a small “cargo recognition α-helix” which is embedded in an IDR. This region undergoes a structural transition from a random coil to an α-helix upon binding to BicD2. In single molecule TIRF assays, four single point mutations within the cargo recognition helix significantly inhibited the interaction between BicD2 and Nup358^min-zip^, and impaired run length and speed of the mutant DDBN^min-zip^ complexes on microtubules. Our results may facilitate the identification of regulatory mechanisms for BicD2 dependent transport pathways, which are important for cell cycle control, brain and muscle development and vesicle transport.

Activation of BicD2 is a key regulatory step for transport, as it is required to activate dynein for processive motility. The cargo recognition α-helix may be a structural feature that stabilizes BicD2 in its activated state. We propose that binding of cargo induces a coiled-coil registry shift in BicD2, which promotes loop-opening and activates BicD2 for dynein/dynactin binding. Notably, Nup358 interacts more strongly with the dynein/dynactin/BicD2 complex compared to BicD2 alone, which supports the loop-opening mechanism for activation. We also show that BicD2 and KLC2 bind to spatially close, but non-overlapping binding sites on Nup358, supporting the hypothesis that Nup358 is capable of simultaneously recruiting dynein and kinesin-1 machineries to the nucleus for bi-directional transport.

## Methods

### Protein expression and purification

Nup358-min and BicD2-CTD were expressed and purified as previously described^30,50,59^.

For details, see SI.

### Pull-down assays

GST-pull down assays of human Nup358-min-GST and human BicD2-CTD were performed as described^59^. For details, see SI.

### Nuclear Magnetic Resonance (NMR)

HSQC NMR experiments of ^15^N-labeled Nup358-min were recorded on a 0.2 mM 440 μl sample with 10% D_2_O on a Bruker 800 MHz spectrometer equipped with a cryoprobe at 25 °C. HSQC was taken on a 1:1 mixture of ^15^N-Nup358^min^-BicD2-CTD, where the sample was concentrated to keep the concentration of Nup358-Min to 0.2 mM in 20 mM HEPES pH 7.5, 150 mM NaCl, 0.5 mM TCEP. For backbone assignment were accomplished using standard triple resonance experiments, and standard NMR processing analysis tools^79,80,81,82^. For details see SI. ^15^N CEST NMR was initially performed with a 0.6 mM sample and a 10:1 ratio of BicD2-CTD:^15^N-Nup358-min. The temperature was reduced to 20 °C, and the pH was reduced to 6.5 to help improve signal/noise by decreasing the rate of solvent exchange of amide protons. ^15^N DCEST NMR^83,84^ was performed with the same sample conditions. A second sample with a 20:1 ratio of 15N-Nup358-min: BicD2-CTD was utilized to improve the S/N ratio of the weakest peaks. For both experiments, the 800 MHz spectrometer with cryoprobe was used. For CEST, the saturation pulse was 400 ms at both 10 Hz and 20 Hz. The saturation frequency ranged from 118 ppm (the center of the HSQC spectrum) to 126 ppm in steps of 0.25 ppm in CEST. The second attempt was run from 116 to 124 ppm. The DCEST was performed to cover the whole spectral width for amides, with a 10 Hz saturation and a 700 Hz spectral width, and a 20 Hz saturation pulse with a 600 Hz spectral width. ^13^C’ HNCO-CEST NMR^56^ was performed with a 0.6 mM sample and a 20:1 ratio of BicD2-CTD:Nup358-Min at 20 °C and pH of 6.5. For this experiment, a 600 MHz spectrometer with cryoprobe was used to minimize the effect of^13^C’ transverse relaxation. The saturation pulse was 300 ms at 10 Hz. The saturation frequency was ranged from 170 to 180 ppm in steps of 0.33 ppm. The CEST data were fitted with the program RING NMR Dynamics^57^ and are presented in Tables 1 and S1.

### CD spectroscopy

CD spectroscopy was performed as previously described^59^. For details, see SI.

### SAXS experiments

Nup358-min and BicD2-CTD were purified as described above. The monodispersity of the protein was confirmed by SEC-MALS which is published^47,50^. Purified Nup358-min and BicD2-CTD was dialyzed against the following buffer: 150 mM NaCl, 20 mM HEPES pH 7.5, 0.5 mM TCEP. The dialysis buffer was used as buffer match for SAXS experiments as well as for dilutions. The following protein concentrations were used: Nup358-min 4 mg/ml, BicD2-CTD 1 mg/ml. To assemble the complex, Nup358-min and BicD2-CTD were mixed in 1:1 molar ratio and incubated for 30 minutes on ice, with a final concentration of 1.3 mg BicD2-CTD and 1.3 mg Nup358-min. The affinity of BicD2-CTD towards Nup358-min is 1.7 ±0.9 μM (Fig. 1), therefore this protein concentration is sufficient for complex formation. To assure monodispersity, SAXS-data were collected for at least 3 protein concentrations for each sample, and we also collected data of a Nup358-min/BicD2-CTD complex that was further purified by gel filtration, with comparable results (data not shown). Prior to data collection, samples were thawed, filtered (pore size 0.2 μm) and centrifuged (30 min, 21,700 g, 4°C).

SAXS data was collected at the beam line 7A1 at the Cornell High Energy Synchrotron Source (CHESS), with a dual Pilatus 100k detector system (Dectris, Baden, Switzerland), at a single detector position, on July 3, 2019 as described previously^85^. Quartz capillary with a path length of 1.6 mm was used as the sample cell (OD=1.5 mm, wall thickness=10 μm). For each dataset, 20 frames were collected at 4°C, with 0.1 s exposure times (wavelength = 9.835 keV, beam dimensions = 250 * 250 μm, beam current = 49.9 mA (positrons), beam flux = 2.4*10^12^ photons/s). Most samples showed no detectable radiation damage, which was monitored by averaging 20 frames.

SAXS data were processed with the BioXTAS RAW software suite (version 2.0.3)^86^. To obtain scattering intensity profiles, 20 data frames were reduced to scattering intensity profiles, placed on an absolute scale, averaged, and the scattering intensity profile of the buffer match was subtracted. The data quality was assessed by Guinier plots, molar mass calculations and dimensionless Kratky plots in BioXTAS RAW^86–90^. Pair distance distribution p(r) functions were derived from the scattering intensity profiles by the program GNOM^91^ of the ATSAS 3.0.0-1 software suite^92^ (implemented in RAW^86^). Fifteen bead model 3D reconstructions were performed with the Dammiff program^93^, (implemented in ATSAS/RAW^86,92^. The resulting models were aligned, grouped into clusters, averaged, and the average model was refined in Dammiff^93–95^. Figures of the refined molecular envelopes were created in the program UCSF Chimera (version 1.14)^96^, developed by the Resource for Biocomputing, Visualization, and Informatics at the University of California, San Francisco, with support from NIH P41-GM103311. P(r) functions were normalized to the highest signal of each curve.

### Protein expression and purification for single-molecule assays

Cytoplasmic dynein and dynactin were purified from 300 gm bovine brain as described in Bingham *et al.*^97^, and tubulin was purified from 200g bovine brain as described in Castoldi and Popov^98^. Purified dynein was stored at −20°C, and dynactin and tubulin were stored at −80°C, in 10 mM imidazole, pH 7.4, 0.2 M NaCl, 1 mM EGTA, 2 mM DTT, 10 μM Mg ATP, 5 μg/mL leupeptin, 50% glycerol. The N-terminal domain of human BicD2 (BicD2^CC1^) with a biotin tag at its N-terminus was expressed in bacteria as described^24^. Full-length human wild type BicD2 was expressed in Sf9 cells as described for *Drosophila* BicD^24^. The Bradford reagent (Bio-Rad, USA) was used to measure the protein concentration. To create a fluorescently labelled version of Nup358-min, the expression vector described above was modified to include a SNAP-tag for fluorescent labelling at its C-terminal domain, which is referred to as Nup358^min^. Nup358^min^ was expressed and purified with the N-terminal GST-tag intact as described for Nup358-min^47,50^, using the BL21-DE3-CodonPlus-RIL strain for expression. Nup358^min^ was dimerized using a leucine-zipper (hereafter called Nup358^min-zip^); the leucine zipper sequence was added at the C-terminus before the snap-tag. The sequences of these two constructs are shown in the Supplemental Methods. The SNAP tag on Nup358^min-zip^ and Nup358^min^ was biotinylated with SNAP-biotin substrate (New England BioLabs, MA) as described^99^.

### Single Molecule assay

Dynein, dynactin, BicD2 and Nu358^min-zip^ constructs were diluted into high salt buffer (30 mM HEPES pH 7.4, 300 mM potassium acetate, 2 mM magnesium acetate, 1 mM EGTA, 20 mM DTT) and clarified for 20 min at 400,000 x g to remove aggregates. To form the dynein-dynactin-BicD2-Nup358^min-zip^ (DDBN^min-zip^) complex, BicD2 and Nup358^min-zip^ were mixed with 565nm and 655nm streptavidin Quantum Dots (Qdots) (Invitrogen, CA), respectively, at a 1:1 molar ratio in separate tubes and incubated 15 min on ice. To block excess binding sites on streptavidin Qdots, 5μM biotin was added to both tubes. Labeled BicD2 and Nup358^min-zip^ were then mixed with preformed dynein-dynactin complex at a molar ratio of 1:1:1:2 (400 nM dynein, 400 nM dynactin, 400 nM BicD2 and 800 nM Nup358^min-zip^) and incubated on ice for 30 min in motility buffer (30 mM HEPES pH 7.4, 150 mM potassium acetate, 2 mM magnesium acetate, 1 mM EGTA, 20 mM DTT). The dynein-dynactin-Nup358^min-zip^ (DDN^min-zip^) complex contained Nup358^min-zip^ that was labeled with a 655 nm Qdot. In the dynein-dynactin-BicD2^CC1^ (DDB^CC1^) complex, BicD2^CC1^ was labeled with a 525 nm streptavidin Qdot. The DDBN^min-zip^, DDN^min-zip^ and DDB^CC1^ complexes were diluted in motility buffer (30 mM HEPES pH 7.4, 150 mM potassium acetate, 2 mM magnesium acetate, 1 mM EGTA, 2 mM MgATP, 20 mM DTT, 8 mg/mL BSA, 0.5 mg/mL kappa-casein, 0.5% pluronic F68, 10 mM paclitaxel and an oxygen scavenger system) to a final concentration of 1.25-2.50 nM dynein for observing motion on microtubules. The oxygen scavenging system consisted of 5.8 mg/ml glucose, 0.045 mg/ml catalase, and 0.067 mg/ml glucose oxidase (Sigma-Aldrich-Aldrich). To analyze the motion of the complexes without any ambiguity, oxygen scavengers were not used in two-color experiments so that the microtubules would photo bleach. Purified tubulin was mixed with rhodamine labeled tubulin at a molar ratio of 10:1 and polymerized as described^24^. PEGylated glass slides were prepared and coated with 0.3 mg/mL rigor kinesin for microtubule attachment as described^24^. After rinsing 2-3 times with motility buffer to remove excess rigor kinesin, MTs were added. Excess microtubules were removed by rinsing with motility buffer. Then dynein protein samples were added to the glass surface. Motion of DDBN^min-zip^, DDN^min-zip^ and DDB^CC1^ were observed using TIRF microscope as described^24^.

### Microscopy and Data Analysis

To detect the motion of Qdot labeled DDBN^min-zip^ and DDB^CC1^ complexes on microtubules, Total Internal Reflection Fluorescence (TIRF) microscopy was used. The TIRF microscope system is operated by the Nikon NIS Elements software and single molecule images were acquired on a Nikon ECLIPSE Ti microscope equipped with through objective type TIRF. The laser lines 488 and 561 nm were used to illuminate 525nm and 655nm Qdots and rhodamine-labeled microtubules. Typically, 600 frames were captured at 200 ms intervals (five frames/s) using two Andor EMCCD cameras (Andor Technology USA, South Windsor, Connecticut). Individual Qdots were tracked using the ImageJ MTrackJ plugin for run length and speed measurements of single complexes^100^. Run length is defined as the total travel distance by individual complexes, and speed was calculated by dividing the run length by the total time. To determine the characteristic run length, data were binned with Sigma plot (Systat Software, Inc) and fit with the equation p(x) =Ae(-x/λ), where p(x) is the relative frequency, x is the travel distance along a microtubule track and A is the amplitude. Speed was reported as mean ± standard deviation (SD), and run length reported as mean ± standard error (SE). Binding frequency was calculated by counting the number of dual colors Qdots per time per μm microtubule. The number of dual color Qdots (DDBN^min-zip^) were counted on 28-32 microtubules (total ~500 μm) in each case. For measuring the formation of DDBN^min-zip^ and BN^min-zip^ complexes containing WT or mutant Nup358^min-zip^, the number of dual color and single color (green or red) Qdots were counted on multiple 512 x 512 pixel fields. The percentage of colocalization was calculated from the total number of dual color Qdots divided by all dual and single color Qdots. Statistical significance for two sets of run length data were determined by the Kolmogorov-Smirnov test, a nonparametric distribution. For speed data comparison, an unpaired t-test was performed. For three or more data sets of run length or speed, statistical significance was calculated using one-way ANOVA followed by Tukey’s post-hoc test. Statistical differences for binding frequency of DDBN^min-zip^ containing WT or mutant Nup358^min-zip^ were determined by one way ANOVA followed by Tukey’s post-hoc test.

## Author Contributions

JMG, HC and MYA contributed equally to this work and should be cited as co-first authors. JMG, JZ, and CW performed the NMR experiments. HC and SRS purified the proteins, and performed CD and SAXS experiments, analyzed data and made figures. XZ purified proteins, performed gel-based pulldown assays, analyzed data and made figures. MYA performed all the TIRF experiments. EWD performed the ITC experiments, analyzed data and made figures. CW and SRS designed experiments and JMG, CW, MYA, KT and SRS analyzed data, wrote and edited the manuscript.

## Acknowledgements

We thank for Fabien Ferrage, Guillaume Bouvignies and Philippe Pelupessy for help with the CEST experiments and Bruce Johnson for help with fitting of the CEST data. We thank Michael Cosgrove for helpful discussions regarding SAXS. CW is funded by NIH grants CA206592 and AG069039. SRS is funded by NIH grant R15 GM128119 and additional funds came from the Chemistry Department and the Research Foundation of SUNY. KMT was funded by NIH grant R35 GM136288. MYA was funded by NIH grant R03 NS114115. The CD instrument was supported by NIGMS grants 1R01GM125853-02S1 and 3R35GM130207-01S1. SAXS data was collected at beamline 7A1, Cornell High Energy Synchrotron source, supported by NSF award DMR-1829070, and by NIH/NIGMS award GM-124166. We thank Qingqiu Huang and Richard Gillilan for user support at the synchrotron source. We also thank Patty Fagnant, Carol Bookwalter and Elena Krementsova for cloning, protein expression and protein purification. The authors declare that they have no conflicting interests.

