## Supplementary Information for "Coil-to-Helix Transition at the Nup358-BicD2 Interface Activates BicD2 for Dynein Recruitment"

**Table of content**

**Items** **Page**
**Fig. S1.** The ITC titration curve of BicD2-CTD with Nup358-min-GST. 2
**Fig. S2.** Fully assigned $^{15}$N-$^{1}$H HSQC NMR spectrum for Nup358-min. 3
**Fig. S3.** $^{15}$N CEST profiles for Nup358 residues showing exchange between coil state and a helical state. 4
**Fig. S4.** $^{13}$C' CEST profiles for Nup358 residues showing exchange between coil state and a helical state. 5
**Fig. S5.** $^{15}$N CEST profiles for Nup358 residues showing NO exchange between coil state and a helical state. 6
**Fig. S6.** $^{13}$C' CEST profiles for Nup358 residues showing NO exchange between coil state and a helical state. 7
**Fig. S7.** Calibration curve for the CD signal, based on a published melting curve of BicD2-CTD. 8
**Fig. S8**. SAXS plots for Nup358-min/BicD2-CTD complex. 9
**Fig. S9.** SAXS plots for Nup358-min. 10
**Fig. S10.** SAXS plots for BicD2-CTD. 11
**Fig. S11.** The pair distance distribution (p(r)) functions of the Nup358-min/BicD2-CTD complex, Nup358-min and BicD2-CTD. 12
**Fig. S12.** SDS-PAGE analyses of pull-down assays to probe the effect of Nup358 mutants on the interaction with BicD2. 13
**Fig. S13.** Nup358 point mutant L2177A does not affect the motion of dynein dynactin complex. 14
**Fig. S14.** Nup358 point mutations that diminish the interaction with BicD2-CTD also diminish formation of the DDBN complex. 15
**Table S1**. Summary of fits of $k_{ex}$, $P_b$, and $\Delta\delta$ for individual CEST curves. 16
**Table S2**. Summary of SAXS data. 17
**Supplemental Movie Legends** 18
**Supplemental Methods** 19

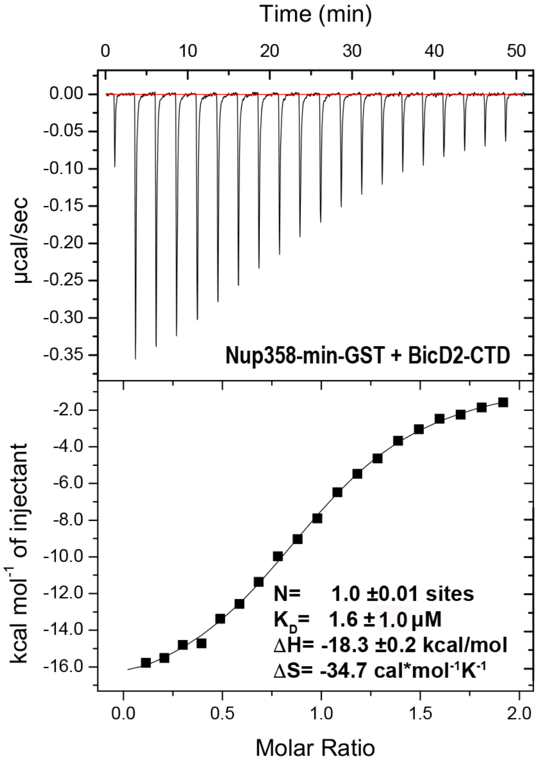


**Fig. S1. The ITC titration curve of BicD2-CTD with Nup358-min-GST** (i.e. with the GST-tag intact) is shown, from which the affinity was determined to be 1.6 ± 1.0 µM, demonstrating that GST fusion has no effect on binding. Related to Fig. 1.

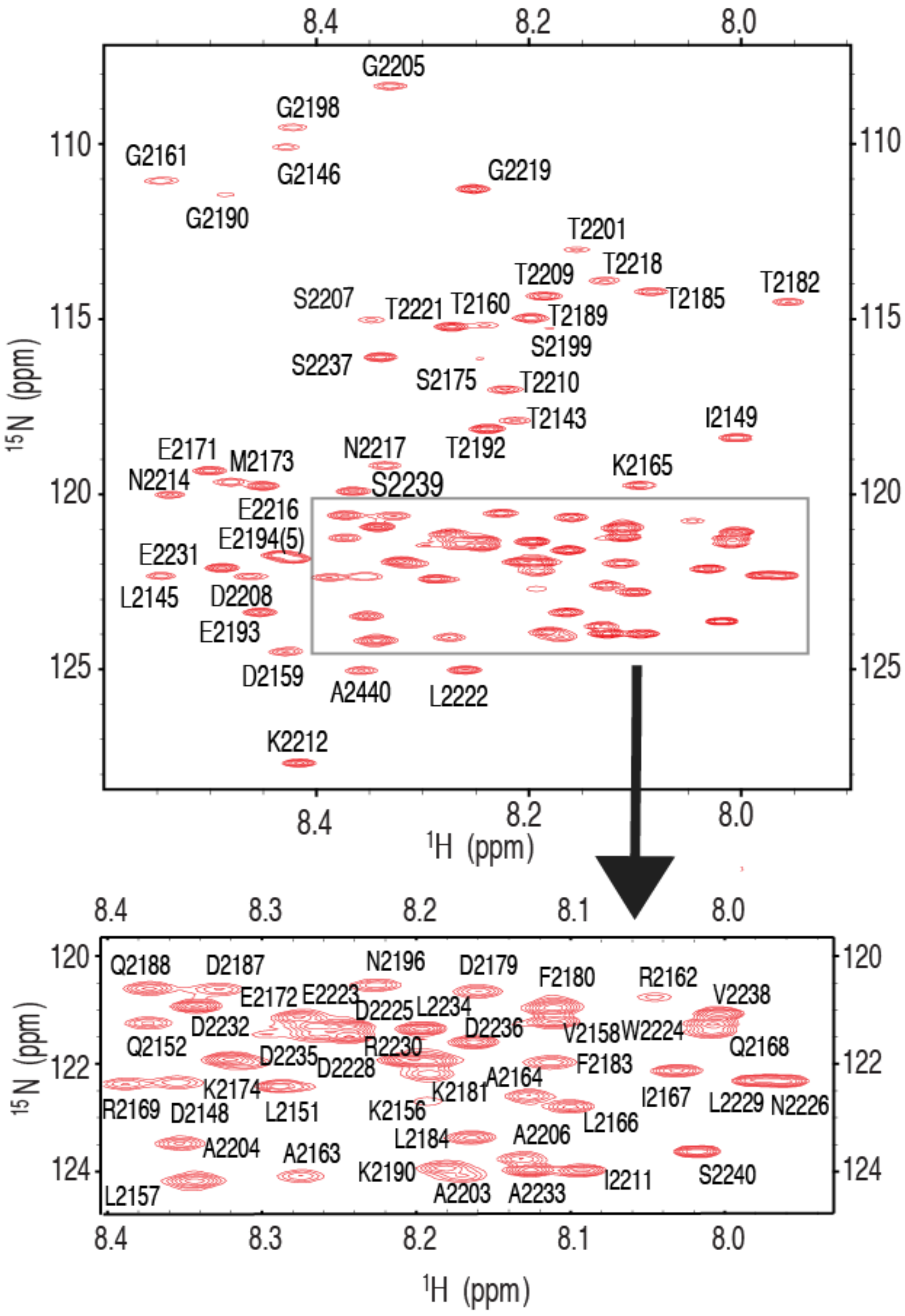


**Fig. S2: Fully Assigned $^{15}$N-$^{1}$H HSQC NMR Spectrum for Nup358-Min.** Inset shows the assignment of peaks in a crowded region.

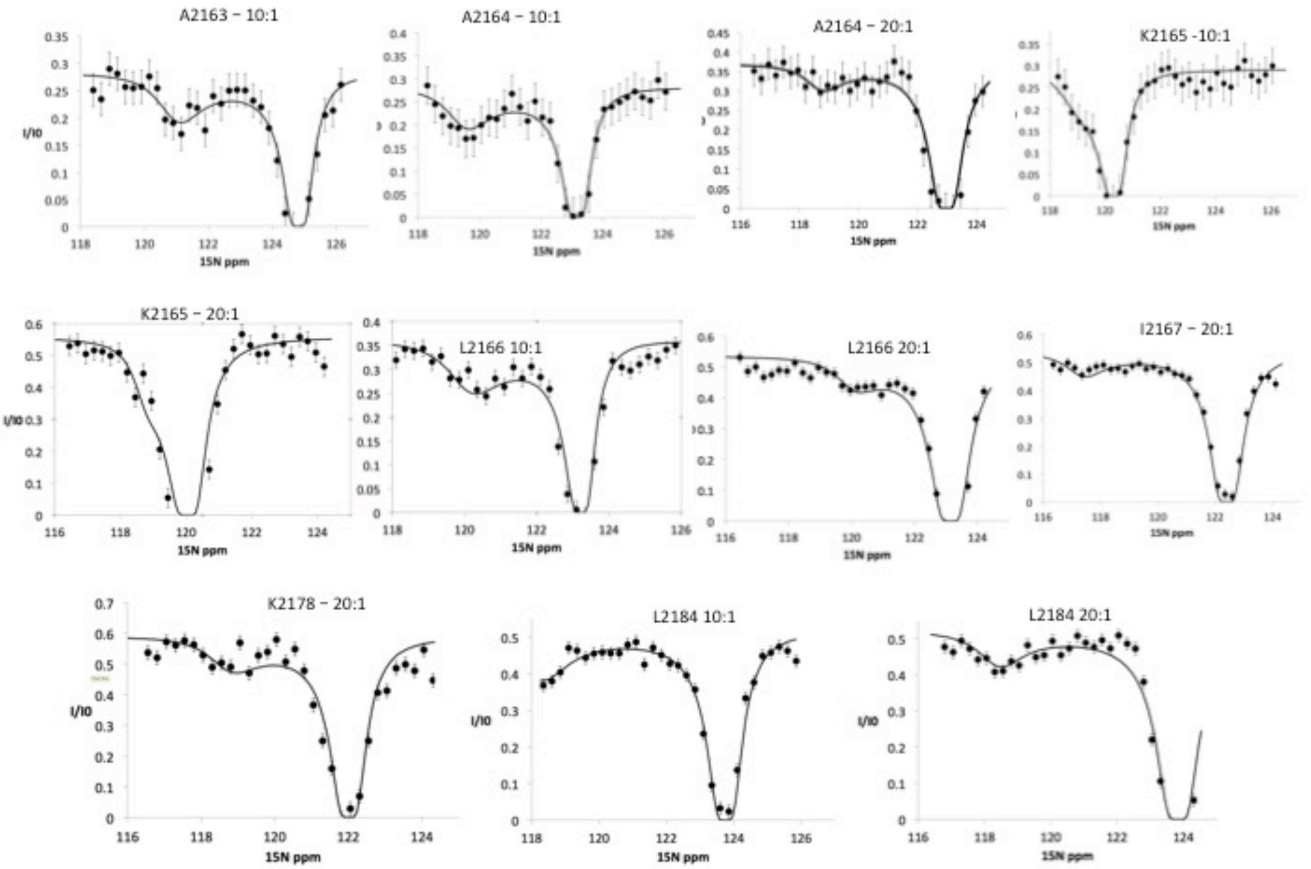


**Fig. S3: $^{15}$N CEST profiles for Nup358 residues showing exchange between coil state and a helical state.** These CEST profiles have a double-dip appearance, showing the bound state, represented by the minor dip as well as the free state, represented by the major dip. This experiment was repeated under slightly different conditions, one with a molar ratio 10:1 Nup358-min to BicD2-CTD and another one with a ratio of 20:1 Nup358-min to BicD2-CTD. The chemical shift range was also shifted by 2 ppm when the 20:1 ratio experimental was run. For example, one can see the presence of both the minor and major dips in L2184 in the “10:1” experiment as well as the “20:1” experiment, although the major dip do not return to baseline. This truncation was due to limited NMR time, necessitated by the stability of the sample. The $^{15}$N CEST curves resulted in the following residues being assigned as part of the $\alpha$-helical region in bound state: A2163, A2164, K2165, L2166, I2167, K2178, and L2184.

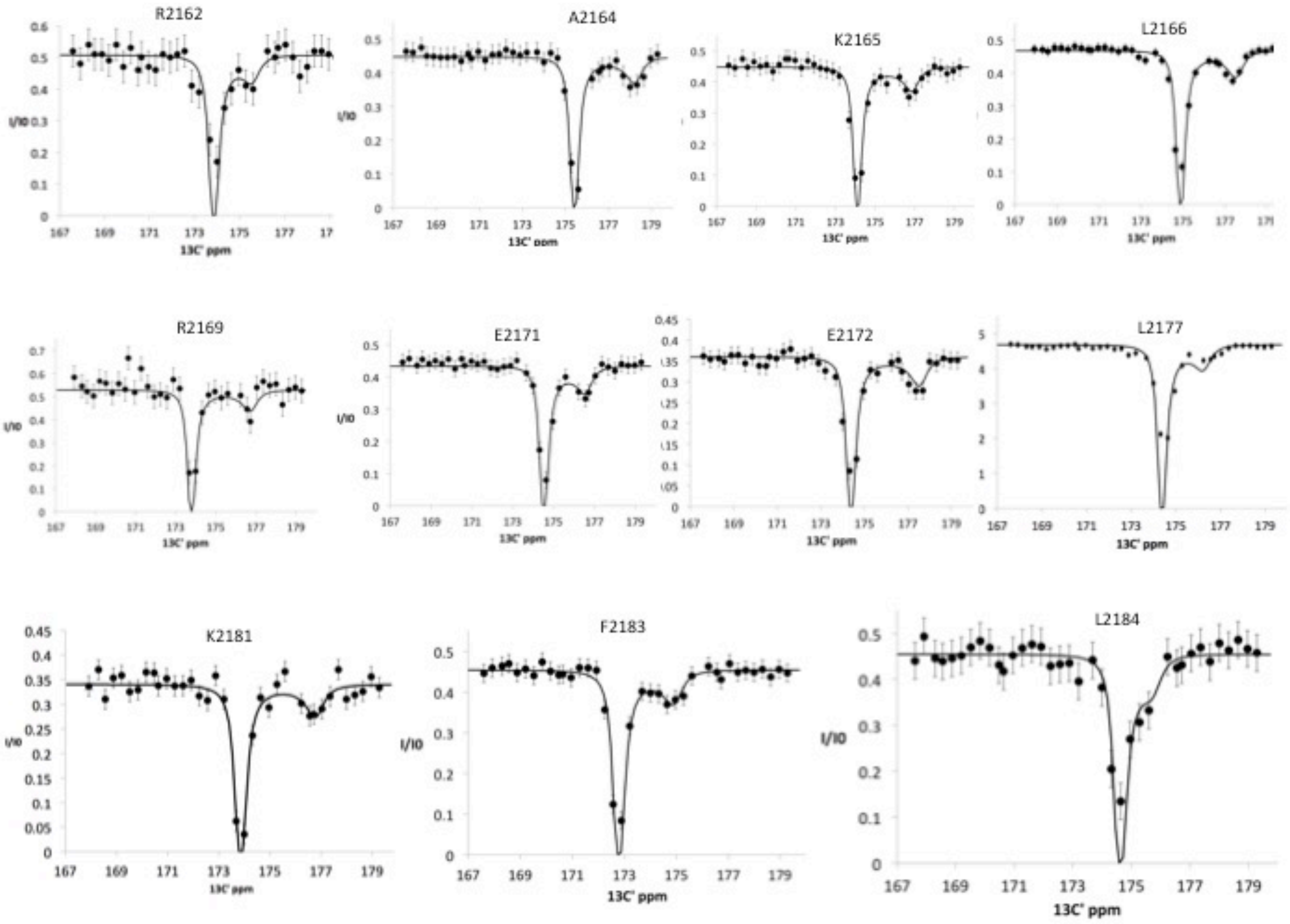


**Fig. S4: $^{13}$C' CEST profiles for Nup358 residues showing exchange between coil state and a helical state**, showing the minor dip as well as the major dip, representing the chemical shift of the bound state. This experiment was performed using a 20:1 molar ratio of Nup358-min to BicD2-CTD. The following residues, already identified as $\alpha$-helical residues, using the $^{15}$N CEST experiments, were confirmed with $^{13}$C' CEST experiments: A2164, K2165 and L2166. Additionally, the following residues were added to the list of $\alpha$-helical residues: R2162, R2169, E2171, E2172, L2177, K2181, F2183, and L2184. To fit the global exchange rate, only curves with sufficient signal/noise, significant resolution between the major and minor dips, and curves with sufficient resolution between the minor dip and the end of the curve were used.

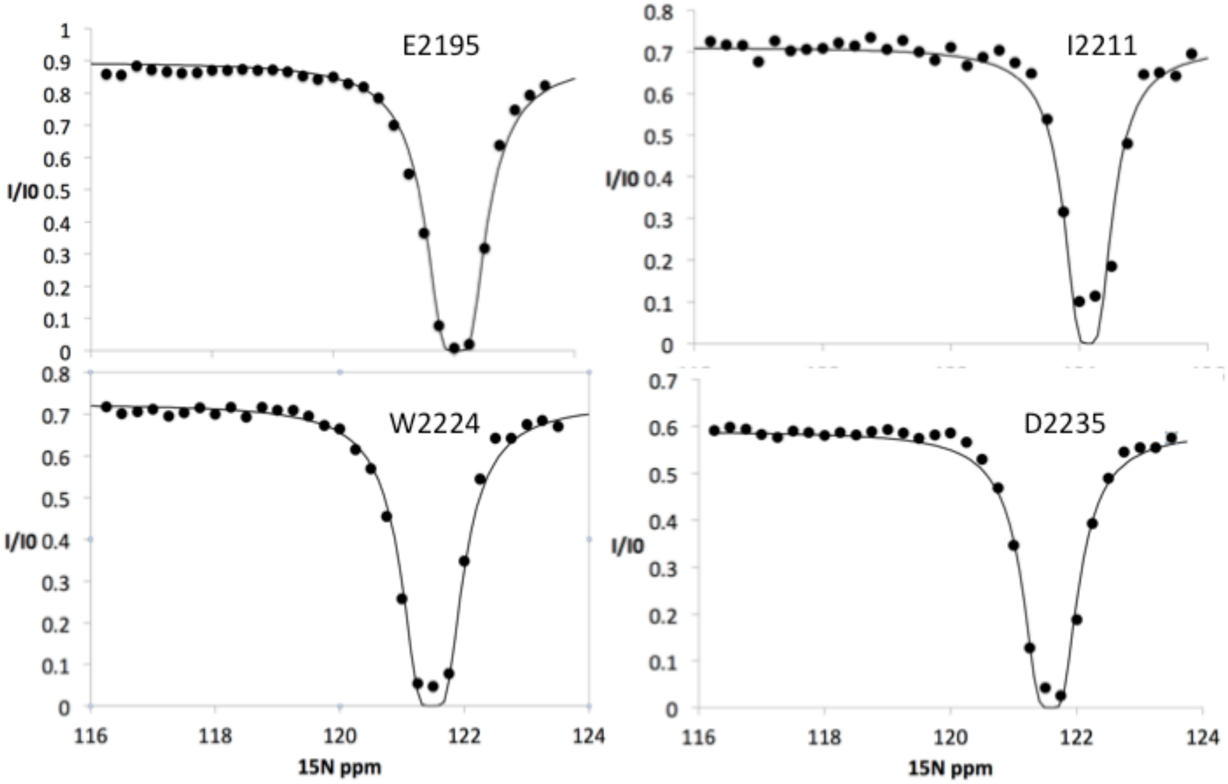


**Fig. S5: [15]N CEST profiles for Nup358 residues showing NO exchange between coil state and a helical state.** These [15]N CEST curves show only a single dip at the major chemical shift, indicating that these sites do not undergo chemical exchange to a bound state. Note that these sites include residues around the LEWD motif as well as sites all the way to the C-terminus of Nup358-min.

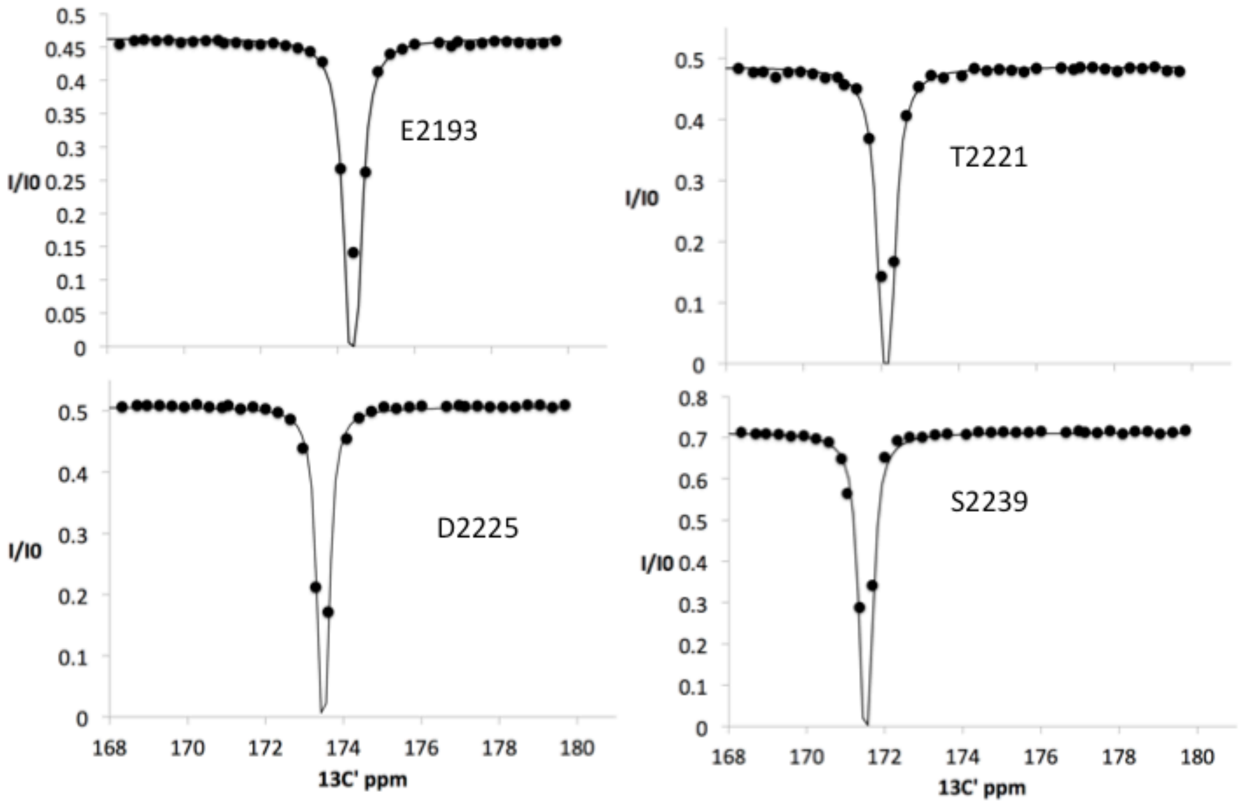


**Fig. S6: $^{13}$C CEST profiles for Nup358 residues showing NO exchange between coil state and a helical state,** showing only a single dip at the major chemical shift. Note that these sites include residues around the LEWD motif as well as sites at the C-terminus half of Nup358-min (see Fig. 4 in the main text).

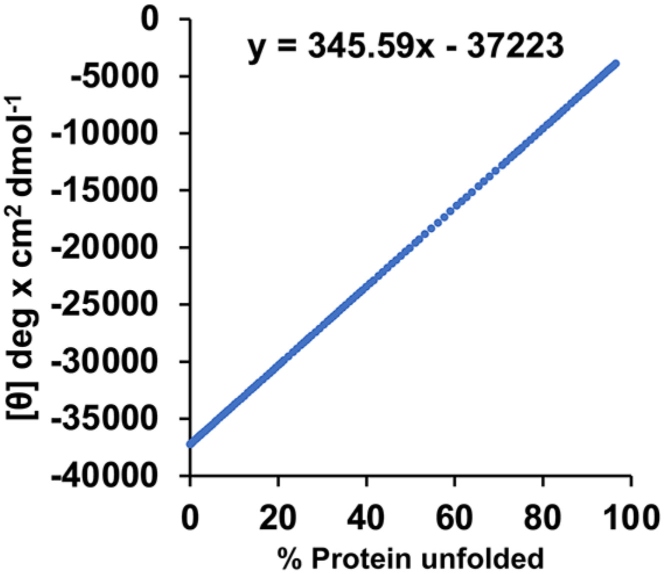


**Fig. S7. Calibration curve for the CD signal, based on a published melting curve of BicD2-CTD**. We previously published the thermal unfolding curve of wild type BicD2-CTD that was recorded by CD spectroscopy at 222 nm (Fig. 4E in reference [30]). Here, this curve was re-plotted as mean residue molar ellipticity [Θ] versus % protein unfolded, to obtain a calibration curve for the α-helical content. 0% and 100% protein unfolded represent the values of $[\Theta]_{min}$ and $[\Theta]_{max}$ from the melting curve respectively. At the apparent melting temperature $T_M$, the protein is 50% unfolded. For purposes of calibration we assume that native, 0% unfolded BicD2-CTD has an α-helical content of 96% as observed in the crystal structure[30]. We also used this method recently to assess changes in α-helical content in the *Drosophila* homolog BicD[47].

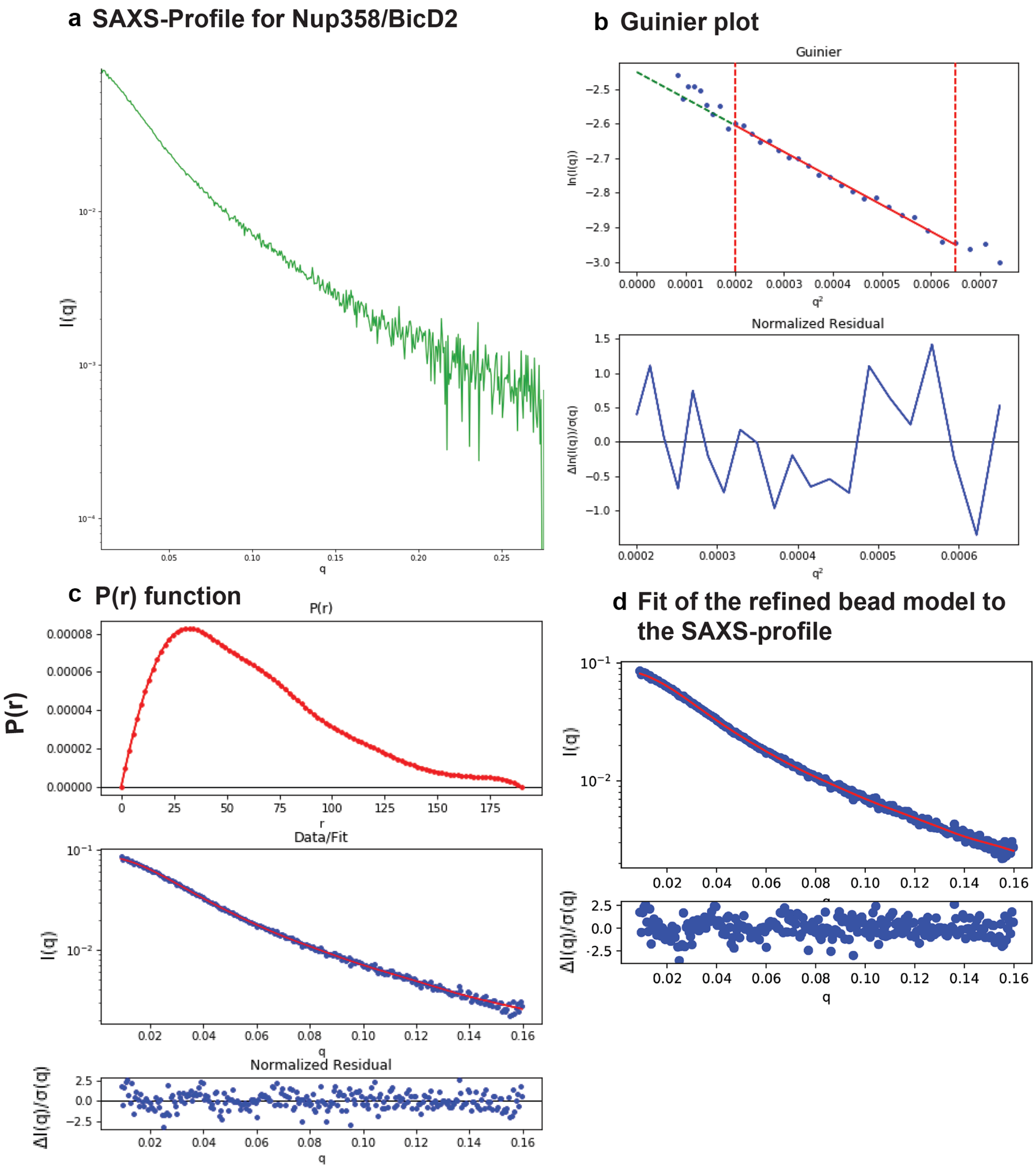


**Fig. S8. SAXS plots for Nup358-min/BicD2-CTD complex**. (a) The scattering intensity profile I (q) is shown as a function of the scattering vector (q). (b) The Guinier plot (top panel) is shown together with the normalized residual of the Guinier fit (bottom panel). (c) The pair distance distribution function p(r) is shown (top panel). The middle panel shows the scattering intensity profile calculated from the p(r) function (red) overlaid with the scattering intensity profile of the actual data (blue). The bottom panel shows the normalized residual of the fit. (d) The "data" scattering intensity profile (blue) is overlaid with the "model" scattering intensity profile that was calculated from the refined bead model 3D reconstruction (red, top panel). The bottom panel shows the normalized residual of the fit.

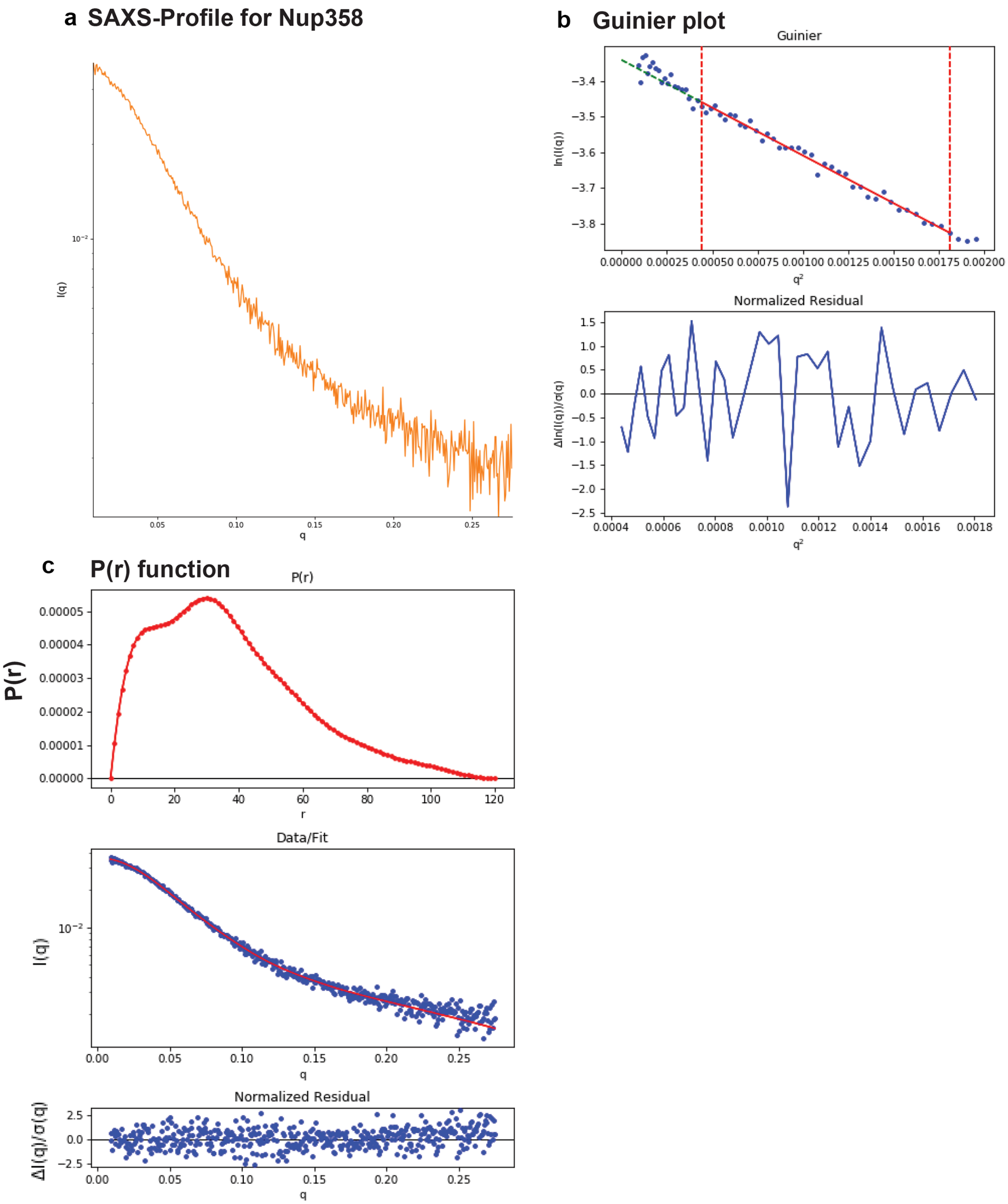


**Fig. S9. SAXS plots for Nup358-min**. (a) The scattering intensity profile I (q) is shown as a function of the scattering vector (q). (b) The Guinier plot (top panel) is shown together with the normalized residual of the Guinier fit (bottom panel). (c) The pair distance distribution function p(r) is shown (top panel). The middle panel shows the scattering intensity profile calculated from the p(r) function (red) overlaid with the scattering intensity profile of the actual data (blue). The bottom panel shows the normalized residual of the fit.

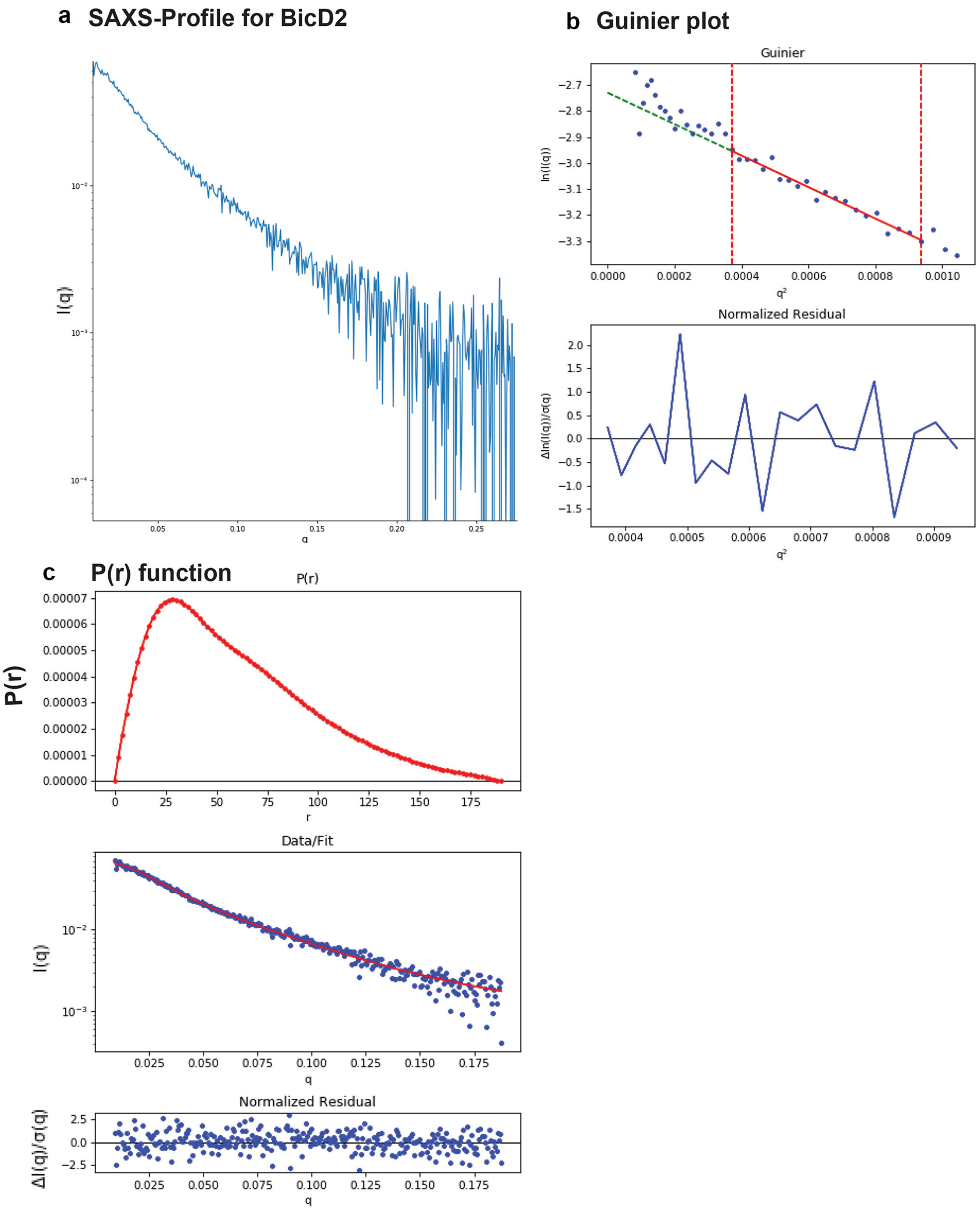


**Fig. S10. SAXS plots for BicD2-CTD**. (a) The scattering intensity profile I (q) is shown as a function of the scattering vector (q). (b) The Guinier plot (top panel) is shown together with the normalized residual of the Guinier fit (bottom panel). (c) The pair distance distribution function p(r) is shown (top panel). The middle panel shows the scattering intensity profile calculated from the p(r) function (red) overlaid with the scattering intensity profile of the actual data (blue). The bottom panel shows the normalized residual of the fit.

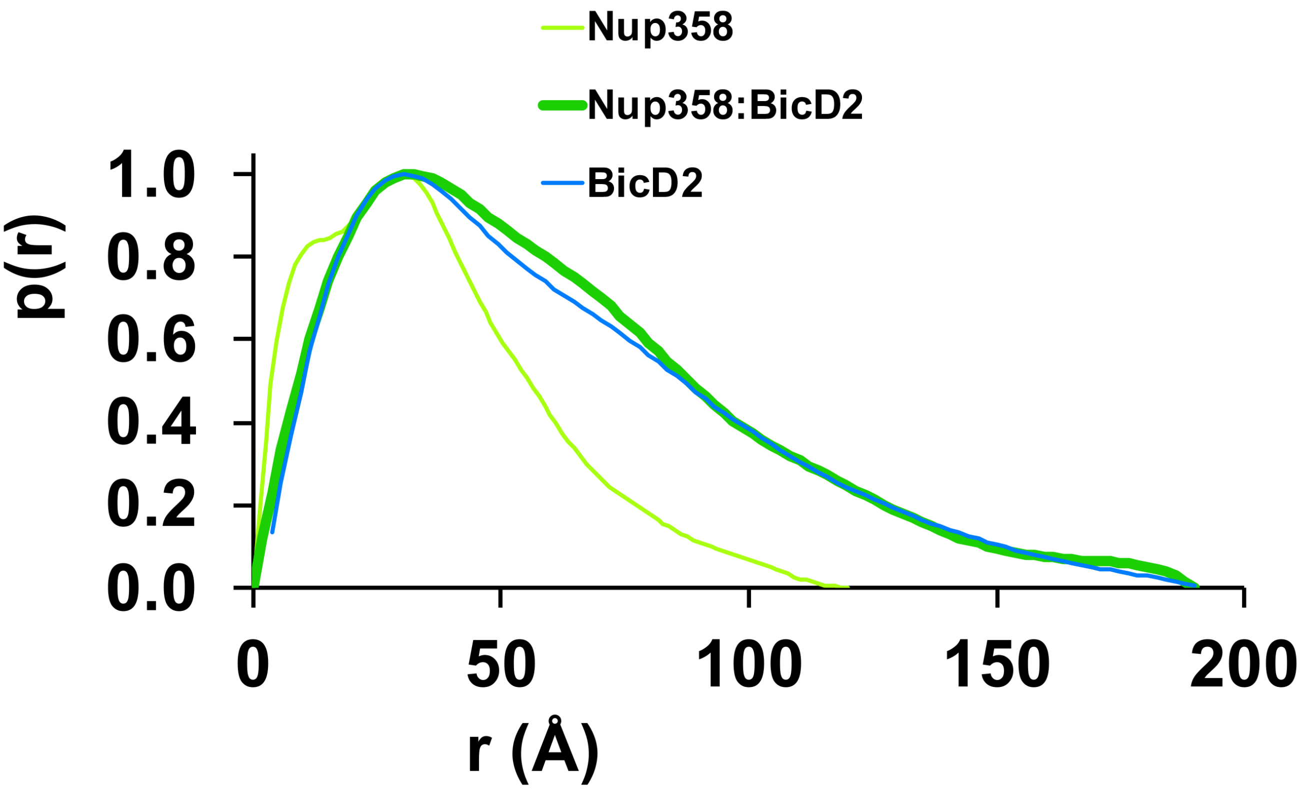


**Fig. S11. The pair distance distribution (p(r)) functions of the Nup358-min/BicD2-CTD complex, Nup358-min and BicD2-CTD** were derived from the SAXS-profiles (Fig. S9 - Fig. S11) and normalized towards the highest signal.

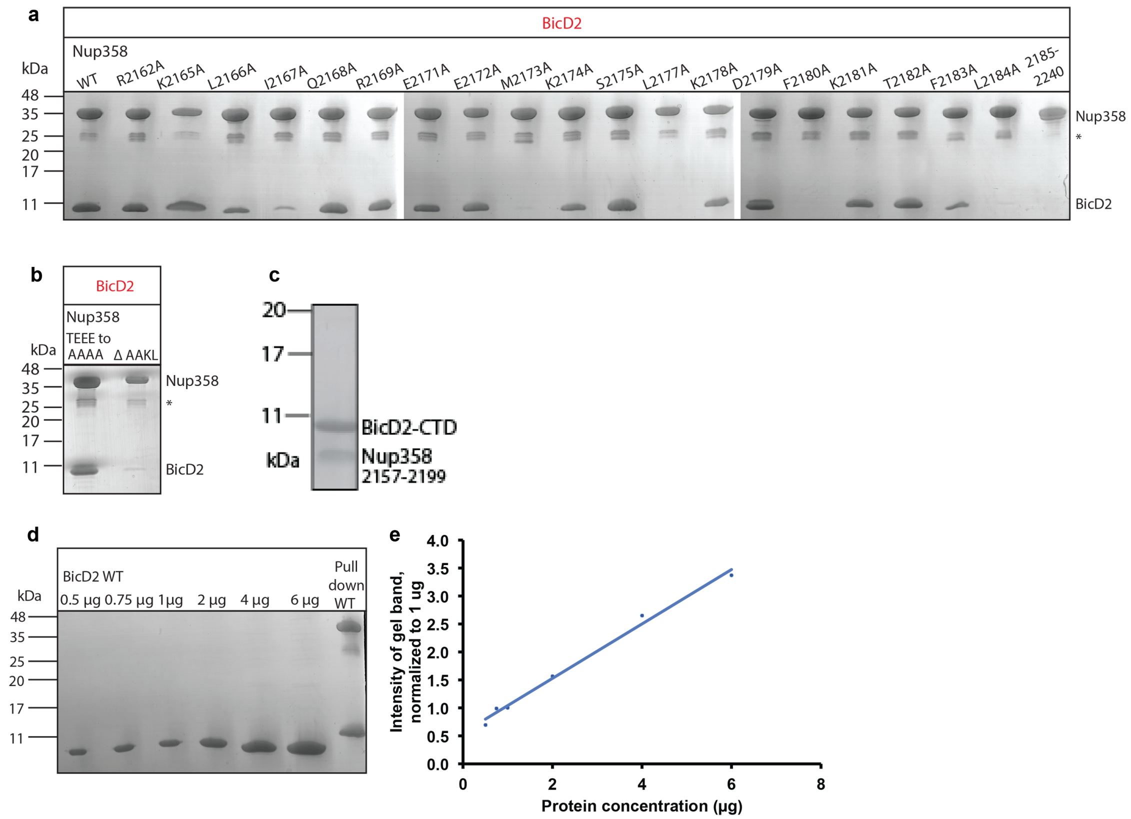


**Fig. S12. SDS-PAGE analyses of elution fractions of the pull-down assays to probe the effect of Nup358 mutants on the interaction with BicD2**. The molecular weights and band position of standard proteins are indicated on the left. (a) A representative result is shown for all mutants that were quantified. Experiments were repeated three times. Note that a small gel band at 25 kDa represents GST. (b) Pull-down of BicD2-CTD and mutants of Nup358-min. Left panel: the linear sequence motif TEEE which is located outside of the cargo recognition helix was mutated to AAAA, to exclude the possibility that it is engages in the interaction. Right panel: The sequence motif AAKL was removed to assess the role of the two alanines for the interaction. All four residues showed large structural changes in CEST. (c) SDS-PAGE analysis of the purified Nup358(residues 2157-2199)/BicD2-CTD complex. The individually purified proteins were mixed and the complex was isolated by gel filtration chromatography. (d) SDS-PAGE analyses of purified BicD2-CTD at known protein concentrations. (e) The signals of the protein bands from c were quantified and the intensities of the gel bands were normalized to the 1 mg gel band. The normalized intensities are plotted versus the protein concentration. Note that the curve is linear.

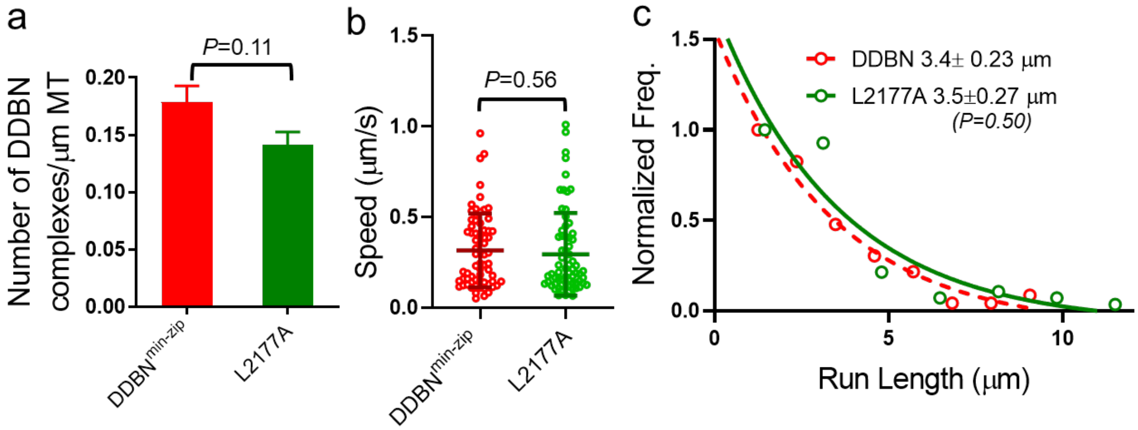


**Fig. S13. Nup358 point mutant L2177A does not affect the motion of dynein dynactin complex.** (a) Histogram showed that the binding affinity of Nup358min-zip-L2177A (0.14±0.012/µm, n=30; taken from Fig. 8a) is similar to WT DDBN$^{min-zip}$ (0.18±0.014/µm (p=0.11, test-test). (b) Nup358$^{min-zip}$ mutant L2177A (green) (0.29 ±0.22 µm; N=69) has the same speed as DDBN$^{min-zip}$ formed with WT-Nup358min-zip (0.32 ±0.20 µm, N=68; red; from Fig. 2h) (p=0.56, t- test). (c) Run length of DDBN formed with WT-Nup358$^{min-zip}$ (red dashed line, from Fig. 2i) is the same as DDBN$^{min-zip}$ formed with Nup358$^{min-zip}$-L2177A (p=0.5, t- test).

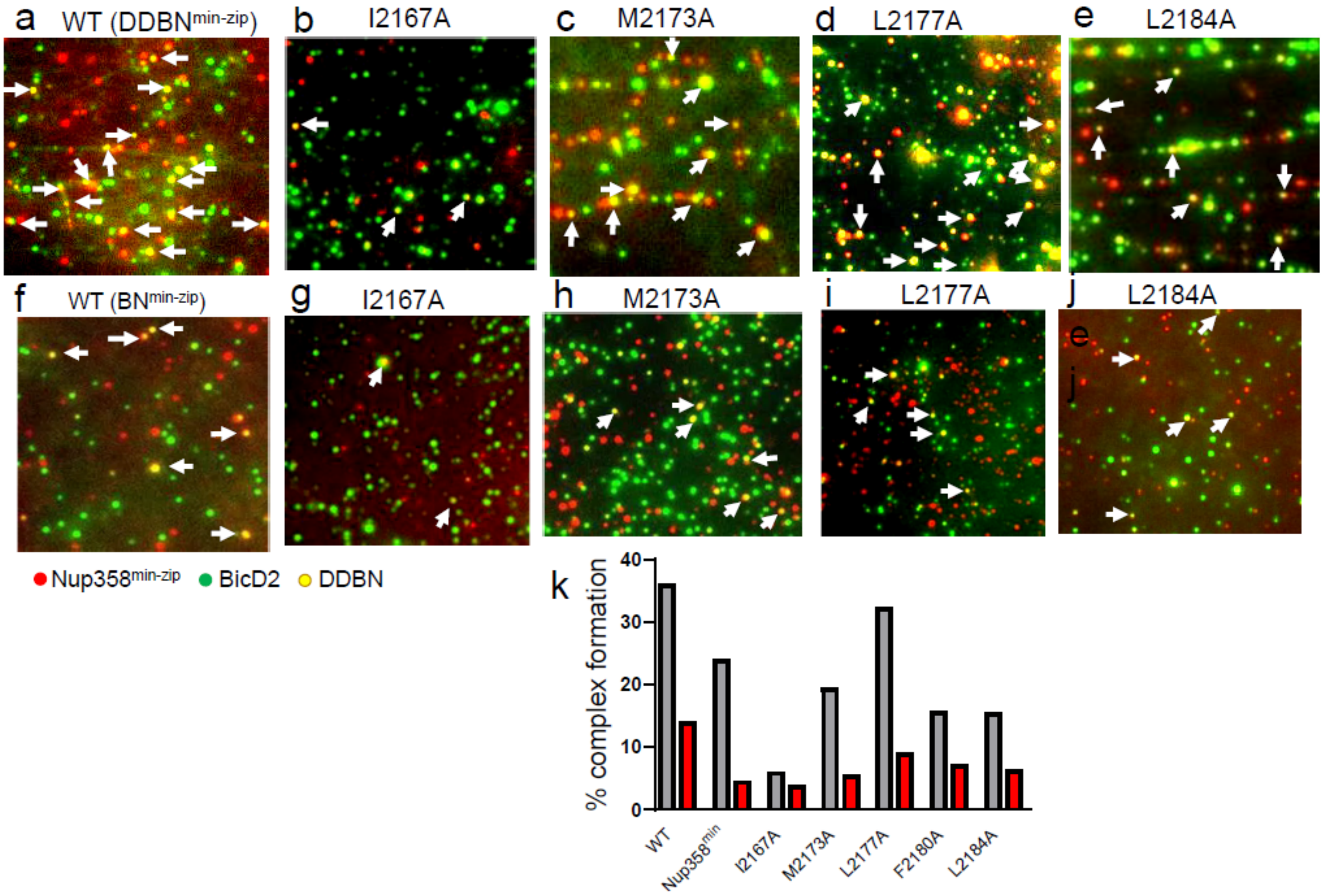


**Fig. S14. Nup358 point mutations that diminish the interaction with BicD2-CTD also diminish formation of the DDBN$^{min\text{-}zip}$ complex.** (a-e) Visualization of complex formation of DDBN$^{min\text{-}zip}$, formed with WT Nup358$^{min\text{-}zip}$ and Nup358$^{min\text{-}zip}$ mutants I2167A, M2173A, L2177A and L2184. DDBN$^{min\text{-}zip}$ formed with mutant F2180A is not shown because of its similarity to L2184. Yellow dots (some indicated by arrows) indicate the formation of the DDBN$^{min\text{-}zip}$ complex, but single color (green or red) Qdots show individual uncomplexed BicD2 and Nup358$^{min\text{-}zip}$ molecules, respectively. (f-j). Visualization of complex formation of BN$^{min\text{-}zip}$ formed with WT Nup358$^{min\text{-}zip}$ and Nup358$^{min\text{-}zip}$ mutants I2167A, M2173A, L2177A and L2184. (k) The dynein/dynactin complex (DDBN$^{min\text{-}zip}$ in gray) enhances the formation of BicD2/Nup358$^{min\text{-}zip}$ complexes (BN$^{min\text{-}zip}$ in red). Note that WT denotes DDBN$^{min\text{-}zip}$ (gray) and BN$^{min\text{-}zip}$ (red), whereas Nup358$^{min}$ denotes DDBN$^{min}$ and BN$^{min}$. The percent formation of BN was: 14.13% (n=580) for WT Nup358$^{min\text{-}zip}$, 4.6% (n=645) for Nup358$^{min}$, 3.9% (n=1142) for I2167A, 5.54% (n=1877) for M2173A, 9% (n=1744) for L2177A, 7.2%(n=1032) for F2180A and 6.3% (n=412) for L2184A Nup358$^{min\text{-}zip}$ . In contrast, these values for DDBN complexes were 36.1% (n=945), 24.1% (n=514) ,6% (n=986), 19.6% (n=347), 32.3% (n=904), 16.7%(n=372) and 15.5% (n=890) respectively. (Figs. 8b and S2a were combined).

**Table S1a. Summary of $^{15}$N CEST Curve Fits at 20:1 Molar Ratio of BicD2: $^{15}$N-Nup358**

| Residue | $k_{ex}$ ($s^{-1}$) | $P_b$ | $\Delta\delta$(ppm) |
|---|---|---|---|
| A2164 | 704± 211 | 0.07±0.02 | -4.3± 0.4 |
| K2165 | 139± 162 | 0.25± 0.08 | -0.8± 0.3 |
| L2166 | 381± 182 | 0.07±0.03 | -3.8± 0.3 |
| I2167 | 975± 624 | 0.02±0.04 | -5.7± 0.2 |
| K2178 | 31± 35 | 0.01±0.01 | -3.6± 0.2 |
| L2184 | 160± 92 | 0.07±0.03 | -5.5± 0.2 |

**Table S1b. Summary of $^{15}$N CEST Curve Fits at 10:1 Molar Ratio of BicD2: $^{15}$N-Nup358**

| Residue | $k_{ex}$ ($s^{-1}$) | $P_b$ | $\Delta\delta$ (ppm) |
|---|---|---|---|
| A2163 | 12± 73 | 0.07±0.05 | -3.6± 0.2 |
| A2164 | 228± 72 | 0.09± 0.02 | -3.6± 0.3 |
| K2165 | 757± 273 | 0.13±0.07 | -1.3± 0.3 |
| L2166 | 297± 195 | 0.05±0.02 | -3.3± 0.6 |

NOTE: The change in chemical shift in Table 1 was taken from Table S1b for the alanine residues due to a clearer minor peak. For the others, the change in chemical shift in Table 1 was taken from Table S1a. The values used in the weighted averages for $k_{ex}$ and $P_b$ were taken from Table S1a, excluding the K2165 and K2178 with abnormal $P_b$ and significant noise in their CEST curves. Including the K2165 and K2178 values, the weighted average $k_{ex}$ value would be 80 ± 30 $s^{-1}$, and the $P_b$ value would be 0.03 ± 0.01.

**Table S1c. Summary of $^{13}$C' CEST Curve Fits at 20:1 Molar Ratio of BicD2: $^{13}$C, $^{15}$N-Nup358**

| Residue | $k_{ex}$ ($s^{-1}$) | $P_b$ | $\Delta\delta$ (ppm) |
|---|---|---|---|
| R2162 | 313± 328 | 0.05±0.03 | 1.7± 0.3 |
| A2164 | 474± 304 | 0.07± 0.03 | 2.8± 0.2 |
| K2165 | 208± 179 | 0.05±0.07 | 2.7± 0.1 |
| L2166 | 391± 170 | 0.04±0.03 | 2.6± 0.2 |
| R2169 | 140± 125 | 0.06±0.01 | 2.9± 0.2 |
| E2171 | 359± 175 | 0.07±0.03 | 2.2± 0.2 |
| E2172 | 8± 98 | 0.22±0.07 | 3.0± 0.1 |
| L2177 | 540± 241 | 0.03±0.02 | 0.9± 0.3 |
| K2181 | 316± 155 | 0.06±0.02 | 3.0± 0.1 |
| F2183 | 464± 98 | 0.06±0.01 | 3.3± 0.3 |
| L2184 | 238± 165 | 0.08±0.04 | 1.0± 0.2 |

### Table S2a. Summary of SAXS data

| Sample | $R_G$ (Å) Guinier plot | $R_G$ (Å) p(r) function | I(0) p(r) function/ I(0) Guinier plot | TE / $\chi^2$ p(r) function | $D_{Max}$ p(r) function | MW */**/*** (kDa) |
|---|---|---|---|---|---|---|
| Nup358/BicD2 | 48.1 ± 0.6 | 55.5 ± 0.2 | 0.086/0.086 | 0.61/1.04 | 190 | 47.7/46.0/47.6 |
| Nup358-min | 28.4 ± 0.3 | 30.6 ± 0.4 | 0.035/0.036 | 0.69/1.05 | 120 | 12.7/12.3/na |
| BicD2-CTD | 42.6 ± 1.0 | 49.7 ± 0.9 | 0.066/0.070 | 0.67/1.10 | 190 | na/na/38.7 |

$R_G$: radius of gyration. I(0): scattering intensity at zero angle. TE: total estimate. $D_{Max}$: maximum particle diameter. MW: molar mass. *MWs were determined from I(0), using glucose isomerase as a standard[89]. **MWs were determined from I(0) without a molar mass standard[89].*** MWs were determined from the volume of correlation as described by Rambo and Tainer[60]. Note that not all methods can be applied to all samples (na). The error of molar masses determined by SAXS is 10%. The calculated MWs are 10.9 kDa for BicD2-CTD and 10.6 kDa for Nup358-min.

### Table S2b. Statistics of SAXS bead model 3D reconstruction

| Sample | Mean normalized spatial discrepancy score (NSD) | $\chi^2$ (refined model) / $\chi^2$ (representative model) | Resolution (Å) | Ambimeter score |
|---|---|---|---|---|
| Nup358-min/BicD2-CTD | 0.70 ±0.05 | 1.08/1.05 | 41 ±3 | 1.23 |

### SUPPLEMENTAL MOVIE LEGENDS

**Movie S1:** Dynein-dynactin-BicD2-Nup358$^{min\text{-}zip}$ (DDBN$^{min\text{-}zip}$) indicated by dual color Qdot (yellow) moving on a microtubule track (not seen due to photobleaching) at 2mM MgATP. Nup358 $^{min\text{-}zip}$ was labeled with a 655nm Qdot (red), BicD2 was labeled with a 525nm Qdot (green), and dynein-dynactin was unlabeled. For both movies, the image was magnified 2X and displayed 4X real speed. Scale bar 2µm. (a) The dual color DDBN$^{min\text{-}zip}$ complex (yellow) moved ~7.4µm in 16 seconds at a speed of 0.46 µm/s. (b) The dual color DDBN$^{min\text{-}zip}$ complex (yellow) moved ~9.3 µm distance in 26 seconds at a speed of 0.36 µm/s. In some cases, BicD2 is not associated with the complex and DDN$^{min\text{-}zip}$ shows diffusive movement (red dot).

**Movie S2:** Dynein-dynactin-Nup358$^{min\text{-}zip}$ (DDN $^{min\text{-}zip}$) complex (red) diffuses on microtubule tracks (not seen due to photobleaching). Nup358$^{min\text{-}zip}$ was labeled with a 655nm Qdot (red), and dynein-dynactin was unlabeled. Image was magnified 2X and displayed 4X real speed. Scale bar 2µm.

### SUPPLEMENTAL METHODS

#### Sequences of expression constructs

**Nup358$^{min}$**

MW 56.5 kDa

MSPILGYWKIKGLVQPTRLLLEYLEEKYEEHLYERDEGDKWRNKKFELGLEFPNLPYYIDGDVKLTQSMAIIRYIAD
KHNMLGGCPKERAEISMLEGAVLDIRYGVSRIAYSKDFETLKVDFLSKLPEMLKMFEDRLCHKTYLNGDHVTHPDF
MLYDALDVVLYMDPMCLDAFPKLVCFKKRIEAIPQIDKYLKSSKYIAWPLQGWQATFGGgDHPPKSDLEVLFQGPL
GS**DIPLQTPHKLVDTGRAAKLIQRAEEMKSGLKDFKTFLTNDQTKVTEEENKGSGTGAAGASDTTIKPNPENTG**
**PTLEWDNYDLREDALDDSVSS**MSGDKDCEMKRTTLDSPLGKLELSGCEQGLHRIIFLGKGTSAADAVEVPAPAA
VLGGPEPLMQATAWLNAYFHQPEAIEEFPVPALHHPVFQQESFTRQVLWKLLKVVKFGEVISYSHLAALAGNPAA
TAAVKTALSGNPVPILIPCHRVVQGDLDVGGYEGGLAVKEWLLAHEGHRLGKPGLG

**Nup358$^{min-zip}$**

MW 60.3 kDa

MSPILGYWKIKGLVQPTRLLLEYLEEKYEEHLYERDEGDKWRNKKFELGLEFPNLPYYIDGDVKLTQSMAIIRYIAD
KHNMLGGCPKERAEISMLEGAVLDIRYGVSRIAYSKDFETLKVDFLSKLPEMLKMFEDRLCHKTYLNGDHVTHPDF
MLYDALDVVLYMDPMCLDAFPKLVCFKKRIEAIPQIDKYLKSSKYIAWPLQGWQATFGGgDHPPKSDLEVLFQGPL
GS**DIPLQTPHKLVDTGRAAKLIQRAEEMKSGLKDFKTFLTNDQTKVTEEENKGSGTGAAGASDTTIKPNPENTG**
**PTLEWDNYDLREDALDDSVSS**MKQLEDKVEELLSKNYHLENEVARLKKLVGERMSGDKDCEMKRTTLDSPLGKL
ELSGCEQGLHRIIFLGKGTSAADAVEVPAPAAVLGGPEPLMQATAWLNAYFHQPEAIEEFPVPALHHPVFQQESF
TRQVLWKLLKVVKFGEVISYSHLAALAGNPAATAAVKTALSGNPVPILIPCHRVVQGDLDVGGYEGGLAVKEWLLA
HEGHRLGKPGLG

**Nup358-min**

10.5 kDa

GPLGS**DIPLQTPHKLVDTGRAAKLIQRAEEMKSGLKDFKTFLTNDQTKVTEEENKGSGTGAAGASDTTIKPNPE**
**NTGPTLEWDNYDLREDALDDSVSS**

**BicD2-CTD**

10.9 kDa

GSH**MYENEKAMVTETMMKLRNELKALKEDAATFSSLRAMFATRCDEYITQLDEMQRQLAAAEDEKKTLNSLLR**
**MAIQQKLALTQRLELLELDHE**

### Protein expression and purification

Proteins were expressed and purified as previously described[30,50,59]. Codon-optimized expression constructs and point mutants were created by a commercial cloning and gene synthesis service (Genscript). Sequences are listed in the supplemental methods. For purification of isotope-labeled Nup358-min, the previously described expression vector for Nup358-min (which encoded the sequence for residues 2148-2240 of human Nup358 cloned into a pGEX6p1 vector)[30] was expressed in the E. *coli* Rosetta 2(DE3)-pLysS strain as described[30], with the following modifications: M9 minimal media (3 g $KH_2PO_4$, 6.8 g $Na_2HPO_4$, 1 g NaCl, 4 g D-glucose, 1g $NH_4Cl$, 2 mM $MgSO_4$, 0.1 mM $CaCl_2$, 100 μg Ampicillin and 35 μg

Chloramphenicol per liter medium) were used for the expression, which was performed at 37 °C. For expression of $^{15}N$ labeled Nup358-min, unlabeled $NH_4Cl$ was replaced in the medium by 1g $^{15}NH_4Cl$ per liter. For expression of $^{13}C$ and $^{15}N$ labeled Nup358-min, 1 g $^{13}C$ D-Glucose (U-13C6) and 1 g $^{15}NH_4Cl$ was used per 1 L of M9 media instead of the unlabeled compounds. Isotopes were obtained from Cambridge Isotope Laboratories.

Nup358-min was purified using the previously described protocol[47], but the cell pellet from a 6 liter bacteria culture was dissolved in 75 ml of the lysis buffer to which 1 ½ tablets of complete EDTA-free protease inhibitor cocktail tablets were added (Roche). In short, Nup358-min was purified by glutathione affinity chromatography and eluted by proteolytic cleavage on the column with PreScission protease. The protein was further purified by size exclusion chromatography as described[47], using the following buffer: 20 mM HEPES pH 7.5, 150 mM NaCl, 0.5 mM TCEP.

BicD2-CTD was expressed and purified as previously described[30,47,50]. In short, BicD2-CTD was purified by Ni-NTA affinity chromatography, followed by proteolytic cleavage of the His6-tag by thrombin. BicD2-CTD was further purified by a second round of Ni-NTA affinity chromatography, followed by size exclusion chromatography in the same buffer as described above. The Nup358 (residues 2157-2199)/BicD2-CTD complex was purified as described for the Nup358-min/BicD2-CTD complex[30].

**Pull-down assays**

GST-pull down assays of human Nup358-min-GST and human BicD2-CTD were performed as described[59]. For the assays, Nup358-min-GST was immobilized on glutathione sepharose beads and incubated with purified BicD2-CTD prior to elution with glutathione. The elution fractions were analyzed by SDS-PAGE (16% acrylamide gels) and stained with Coomassie Blue. ImageJ was used for quantification of gel bands[101].

**Nuclear Magnetic Resonance (NMR)**

For backbone assignment, triple resonance experiments (HNCO, HNCA, HNCACO, HNCOCA, HNCACB, CBCCACONH, and HNN[102] were performed on double-labeled $^{15}N/^{13}C$ Nup358-min at 0.4 mM on a Bruker 800 MHz spectrometer equipped with a cryoprobe. They were performed with non-uniform sampling (NUS) Processing of the data was performed with NMRPipe[79] and SMILE[80]. Further analysis of the data was performed with nmrfam_Sparky[103], including iPine[81,82].

**CD spectroscopy**

Purified proteins at a concentration of 0.3 mg/ml were dialyzed in the following buffer: 150 mM NaCl, 10 mM Tris pH 8 and 0.2 mM TCEP. Data were recorded with a Jasco J-1100 CD Spectrometer, equipped with a thermoelectric control device. A quartz cuvette with a path length of 0.1cm was used. After the buffer baseline subtraction, CD signals were normalized to the protein concentration and converted to mean residue molar ellipticity [Θ]. Thus, the ellipticity Θ (mdeg) was multiplied with the conversion factor 391.51 $cm^2\ dmol^{-1}$ for all spectra with exception of the Nup358-min spectrum, for which the conversion factor was 362.00 $cm^2\ dmol^{-1}$. For the Nup358 + BicD2 spectrum, the spectra of Nup358-min and BicD2-CTD were added together prior to conversion to [Θ].
